# Cryptic diversity within the *Poecilochirus carabi* mite species complex phoretic on *Nicrophorus burying* beetles: phylogeny, biogeography, and host specificity

**DOI:** 10.1101/2021.05.20.443311

**Authors:** Julia Canitz, Derek S. Sikes, Wayne Knee, Julia Baumann, Petra Haftaro, Nadine Steinmetz, Martin Nave, Anne-Katrin Eggert, Wenbe Hwang, Volker Nehring

## Abstract

Coevolution is often considered a major driver of speciation, but evidence for this claim is not always found because diversity might be cryptic. When morphological divergence is low, molecular data are needed to uncover diversity. A taxon for which this holds true are the mites, which are known for their extensive and often cryptic diversity. We studied mites of the genus *Poecilochirus* that are phoretic on burying beetles (Silphidae: *Nicrophorus*). *Poecilochirus* taxonomy is poorly understood. Most studies on this genus focus on the evolutionary ecology of *Poecilochirus carabi sensu lato*, a complex of at least two biological species. Based on molecular data of 230 specimens from 43 locations worldwide, we identified 24 genetic clusters that may represent species. We estimate that these mites began to diversify during the Paleogene, when the clade containing *P. subterraneus* branched off and the remaining mites diverged into two further clades. One clade resembles *P. monospinosus* and *P. austroasiaticus*. The other clade contains 17 genetic clusters resembling *P. carabi s*.*l*.. Among these are *P. carabi sensu stricto, P. necrophori*, and potentially many additional cryptic species. Our analyses suggest that these clades were formed in the miocene by large-scale geographic separation. Diversification also seems to have happened on a smaller scale, potentially due to adaptation to specific hosts or local abiotic conditions, causing some clusters to specialize on certain beetle species. Our results suggest that biodiversity in this genus was generated by multiple interacting forces shaping the tangled webs of life.

## Introduction

Parasites are known for their high biodiversity (García-Varela et al., 2011; Huyse et al., 2005; Magalhães et al., 2007). In all organisms, geographic separation can lead to allopatric speciation, and ecological factors may cause disruptive selection and lead to reproductive isolation in sympatry (Butlin et al., 2012). Ecological effects are amplified by the interaction of parasites with their hosts: Any divergence in host populations can potentially alter selection on the parasites and lead to divergence and even co-speciation with their hosts (Hoberg et al., 1997). Further, reciprocal selection between hosts and parasites may speed up evolution and lead to higher diversification rates among populations (Paterson et al., 2010; Thompson, 2009; Yoder & Nuismer, 2010). These processes can result in many parasite populations or species each specialized on just one host species or population (García-Varela et al., 2011; Hunter & Rosario, 1988; Huyse et al., 2005; Magalhães et al., 2007; Perotti & Braig, 2009). The diversity of parasites is often cryptic, as they tend to live hidden lives and morphological adaptations to parasitism may be similar for all hosts (García-Varela et al., 2011). Hence, the diversity of parasites is likely underestimated.

The mite genus *Poecilochirus* G. & R. Canestrini, 1882 (Mesostigmata: Parasitidae) consists of 20 morphologically described species (Hyatt, 1980; Perotti & Braig, 2009; Ramaraju & Madanlar, 1998). Several species are phoretic as immatures (deutonymphs). They attach to adult burying beetles (Silphidae: *Nicrophorus*) for dispersal (Farish & Axtell, 1971; Schwarz & Koulianos, 1998). The beetles bury small vertebrate carcasses, on which they raise their offspring. While the parental beetles provide brood care, *Poecilochirus* mites dismount from their host beetles, feed on microorganisms, fly eggs, and sometimes beetle eggs, develop into adults, and reproduce (Brown & Wilson, 1992; Meierhofer et al., 1999; Schwarz & Müller 1992).

Deutonymphal mites disperse from the carcass by attaching to the parental beetles or their adult offspring (Schwarz & Koulianos, 1998; Schwarz & Müller, 1992). At larger carcasses, multiple beetle species can co-occur and mites can switch between host individuals. Mites recognize their main host by olfactory cues and some prefer specific *Nicrophorus* species over others (Korn, 1982; Müller & Schwarz, 1990). If the preferred host species is not available, the mites will mount other host species, but their fitness may be reduced when they reproduce along with the ‘wrong’ host species (Brown & Wilson, 1994; Nehring et al., 2017). Such host switches may counteract host specialization (Thompson, 2009).

There is no evidence that the mites affect host beetle fitness during the phoretic dispersal. However, some observations suggest that mites may affect beetle fitness during the reproduction phase. Depending on the environmental conditions, the mites can either directly reduce beetle brood weight and offspring number (De Gasperin & Kilner, 2015; Nehring et al., 2019; Schedwill et al., 2020), or have positive effects on beetle fitness by helping to fend off other competitors such as blowflies, nematodes, or other beetles (Springett, 1968; Sun et al., 2019; Wilson & Knollenberg, 1987). In any case, whether the mites are parasites or mutualists, they have likely coevolved with the host beetles.

*Poecilochirus* species can be distinguished morphologically based on body size, the presence or absence of setae, the form and pattern of setae and the sternal shield, the morphology of the chelicerae, and genital structures (Baker & Schwarz, 1997; Hyatt, 1980; Ramaraju & Madanlar, 1998; Wise et al., 1988). However, the taxonomy of the genus is poorly understood, at least partly because morphological differences between species are small (Mašán, 1999). The best-studied species is *Poecilochirus carabi*, but it has been shown that this species actually consists of at least two reproductively isolated populations that occur sympatrically in Central Europe and that have been named *P. carabi sensu stricto* and *P. necrophori* (Baker & Schwarz, 1997; Hyatt, 1980; Müller & Schwarz, 1990). Field and laboratory studies indicate that *P. necrophori* is a host specialist primarily found on *Nicrophorus vespillo*, while *P. carabi s*.*s*. is prevalent on at least three different *Nicrophorus* species (*N. vespilloides, N. investigator*, and *N. humator*), but rarely on *N. vespillo* (Schwarz, 1996). Furthermore, two reproductively isolated populations of *P. carabi* are specialized on two sympatric North American *Nicrophorus* species (*N. tomentosus* and *N. orbicollis*) and differ in morphology (Brown, 1989; Brown & Wilson, 1992), but their relationship to the European species is unknown. Mites of these morphologically very similar but reproductively isolated and ecologically well-defined populations are considered to belong to a cryptic species complex, termed *P. carabi sensu lato* (Baker & Schwarz, 1990; Masan, 1999).

One might expect that *P. carabi s*.*l*. consists of more than these four cryptic species, given that the holarctic genus *Nicrophorus* includes more than 60 species (Sikes et al., 2002, 2016). Burying beetles originated in the Cretaceous (99–127 Ma) in Asia/Eurasia, colonized the Western hemisphere and have likely back-migrated to Eurasia more than once (Hatch, 1927; Peck & Anderson, 1985; Sikes & Venables, 2013). Today, most species are restricted to single continents, and at most locations, multiple species occur in sympatry (e.g. Brown & Wilson, 1992; Dekeirsschieter et al., 2011). Only *N. vespilloides* and *N. investigator* occur in both the Western and Eastern hemispheres (Sikes et al., 2008; Sikes et al., 2016). *Nicrophorus* species vary in their habitat preferences. Some species inhabit woodlands and others open areas, and differences in the diel activity and reproductive season have been reported for species on both continents (Anderson 1982, Burke et al., 2020; Esh & Oxbrough, 2021; Majka, 2011; Scott, 1998).

Molecular, phylogenetic, and divergence time analyses have been conducted for *Nicrophorus* (Sikes et al., 2008, 2016; Sikes & Venables, 2013), but are missing for *Poecilochirus* species. The scant genetic data that are available are primarily from broader studies on the diversity and phylogenetics of Mesostigmata or Acari, with many samples only identified to the genus level (Klompen et al., 2007; Rueda-Ramírez et al., 2019; Young et al., 2019). The lack of molecular data on the diversity of this genus prevents further insight into its ecology and evolutionary history.

In this study, we want to understand the evolutionary history of *Poecilochirus* mites that are phoretic on *Nicrophorus* beetles, and *P. carabi s*.*l*. in particular, using molecular analyses. Given the biodiversity and wide distribution of the host genus, and the independent descriptions of reproductively isolated mite populations in Europe and North America, we predicted that genetic diversity within the genus *Poecilochirus* is currently underestimated. Based on the results of previous studies (Brown & Wilson, 1992; Müller & Schwarz, 1990), host specificity is expected to be evident in mite genetics, as much of the mites’ reproductive isolation is suggested to be caused by diverging host preferences. We sequenced three genetic markers of *Poecilochirus* mites collected with their *Nicrophorus* hosts on four different continents. We documented their genetic diversity, reconstructed the phylogenetic relationships, and estimated evolutionary divergence times to better understand the drivers of speciation in this mite clade.

## Materials and Methods

We collected *Poecilochirus* mites from burying beetles from North and South America, Europe and Asia and used nuclear and mitochondrial DNA sequences to identify genetic clusters. We then reconstructed phylogenetic relationships, and analyzed the biogeography and host specificity of the main mite clusters.

### Sampling

We focused on mites from burying beetles that morphologically resemble *P. carabi* in their habitus and body size (body length ca. 1 mm; 218 individuals). Samples originated from 43 different locations ranging from Alaska (USA), through Europe, Central Asia, Japan and Melanesia (Figure S1). They were collected together with their host beetles, including 31 *Nicrophorus* species and one carabid beetle (*Pterostichus melanarius*). Specimens were sampled from the wild between 1998 and 2020, and were preserved either in 96% ethanol, propylene glycol, or kept dry (Supplement Table S1). Several specimens of the German populations (Mooswald, Freiburg) could be identified as *P. necrophori* or *P. carabi s*.*s*. by host preference tests following the description of Nehring et al. (2017).

In addition, specimens identified as *P. subterraneus* (n = 12) and *Macrocheles* sp. (n = 2) were added to the data set to serve as the outgroup in phylogenetic analyses (Supplement Table S1). Mite vouchers are deposited in the Sikes Research Collection at the University of Alaska Fairbanks, the Canadian National Collection of Insects, Arachnids, and Nematodes, and the Acarological Collection at the University of Graz.

### Molecular Methods

We extracted DNA from 232 mites. Two different methods were used for DNA extraction. We applied either the Chloroform/Isopropanol method where the whole individual was ground up in liquid nitrogen, or a non-destructive approach using the DNeasy Blood and Tissue Kit (Qiagen). For the non-destructive method, we incubated the entire specimen in 50µl lysis buffer (ATL buffer) and 10μl Proteinase K for approximately 24h at 56°C. After 6–8h, additional 5µl Proteinase K were added. Subsequently, we removed the undamaged specimens and followed the default protocol of the DNeasy Kit. Mite voucher specimens following DNA extraction were preserved in absolute ethanol.

A partial sequence of the Cytochrome Oxidase I gene (COI) was amplified using the universal primer pair LCO1490 and HCO2198 (Folmer et al., 1994). Furthermore, the ribosomal Internal Transcribed Spacer gene (ITS) was amplified. The primer pair used for the ITS amplification was based on the sequences published by Navajas et al. (1999), but we modified them slightly for better annealing (forward: 5’-AGTCGTAACAAGGTTTCCGTAG-3’; reverse: 5’-GGGGGTAATCGCACTTGA TTTC-3’). Additionally, the gene coding for the Large Subunit of the rRNA (LSU) was amplified partly using the universal primer pair LSU-D1D2.fw2 and LSU-D1D2.rev2 (Sonnenberg et al., 2007). Amplification was performed with 30 thermocycler cycles. PCR products were purified either by a polyethylene glycol (PEG)-based method or enzymatically with ExoSAP-IT™ (ThermoFisher). We sent PCR products to Macrogen Europe Inc. (Amsterdam, The Netherlands) for forward and/or reverse Sanger sequencing.

### Identification of genetic clusters

The COI, ITS, and/or LSU sequences of 230 *Poecilochirus* samples were aligned gene-wise using default settings of the Geneious Prime implementation MAFFT v.7.450 (Katoh & Standley, 2013). Alignments were concatenated to a supermatrix in which sequences of “N” symbolized missing data/genes. The supermatrix served as input for a phylogenetic analysis with IQtree multicore version 1.6.12 (Nguyen et al., 2015). IQtree estimates maximum-likelihood (ML) phylogenies using a fast and effective stochastic search algorithm. We used IQtree’s model finder (Kalyaanamoorthy et al., 2017) to select accurately best-fitting evolutionary models for each gene of the supermatrix. The models TPM2u+F+I+G4, TIM3+F+G4, and HKY+F+G4 were chosen for the COI, ITS, and LSU gene block, respectively. The phylogenetic analysis ran with 10,000 bootstrap replicates using the ultra-fast bootstrap approximation for branch support (Minh et al., 2013), and with a parametric approximate likelihood-ratio test (SH-aLTR) testing the null hypothesis that assumes inferred branches with a length of 0 (Anisimova et al., 2011). We set *P. subterraneus* as outgroup. The phylogenetic tree was visualized with FigTree version 1.4.4 and edited using Inkscape 1.0.1. We also used the “Poisson Tree Process” (PTP) model implemented in the tool multi-rate (m)PTP version 0.2.4 (Kapli et al., 2017). The model is a tree-based approach for species delimitation. It suggests the most likely number of species based on maximum likelihood and a Markov Chain Monte Carlo (MCMC) algorithm to provide support values for each putative species clade. Initially, the minimum branch length was calculated (--minbr_auto) based on the input alignment. In a second step the resulting values and the phylogenetic tree were input to the main mPTP analysis with the following parameters: --mcmc 100,000,000; -- mcmc_sample 10,000; --mcmc_burnin 500,000; --mcmc_runs 4; --mcmc_startrandom. To illustrate the geographic distribution of the identified genetic clusters, the R packages maptools v1.0-2 and scatterpie v0.1.5 (R Core Team, 2020) were used to plot the relative abundance of the different clusters at each locality on a world map. To support cluster delineation, uncorrected p-distances within and between clusters were calculated based on the COI alignment using MEGA version 10.1.7 (Kumar, Stecher, Li, Knyaz & Tamura, 2018).

### Morphological identification

Mite specimens that were not destroyed during DNA extraction (n = 95) were mounted and clarified in Heinze-PVA medium and stored in an oven at 50°C until total clarification. Morphological and morphometric analyses of mites were performed using differential interference contrast in a compound microscope (Reichert Diavar, Vienna). Identification of mites was based on the key by Hyatt (1980) and the description of *P. monospinosus* by Wise et al. (1988).

### Host specificity

Host specificity was calculated as Shannon-Wiener Diversity (H’) and Evenness. We performed these calculations for three European and three North American clusters that contained enough samples and host species. In addition, we used a χ^2^-test to investigate if the frequency of host species occupied by a mite cluster deviates from the overall host species frequency in the same geographical area. The host species frequency was derived from the number of mites that we sequenced from each beetle species; whenever possible, we had selected mites from different beetle individuals. Subsequently, the χ^2^ value of each cluster (χ^2^(_cluster_)) was set in relation to the theoretical χ^2^ maximum of the respective cluster (χ^2^(_max_)). A high quotient of χ^2^(_cluster_)/χ^2^(_max_) suggests host specificity of the mite clusters for that area. Calculations for the Shannon-Wiener Index and Evenness were performed using the open source spreadsheet tool provided by LibreOffice version 6.2.8.2., and χ^2^ values were calculated with the R Stats package v3.6.2.

### Phylogenetic inference

We used 38 samples for which sequences of all three genes were available for phylogenetic analyses. These samples covered 16 of the clusters previously identified. We applied Maximum Likelihood (ML) and Bayesian Inference (BI) approaches and used four different methods for assigning confidence levels to branches - SH-aLRT, ultrafast bootstrapping (UFBoot), standard bootstrapping (SBS), and posterior probability (PP). Phylogenetic analyses were carried out with IQTree (ML; aLRT/UFBoot), RaxML version 8.2.4 (ML; SBS; Stamatakis, 2014), and MrBayes 3.2.7a (BI; PP; Ronquist et al., 2012). The input for all analyses is a concatenated alignment generated with Geneious Prime. All analyses were conducted with the data partitioned by gene. Best-fitting substitution models were found using IQtree. Models were adjusted to the most similar substitution model RaxML and MrBayes can integrate. For the IQtree analysis the same parameter settings were used as described above. For the RaxML analysis, we chose the rapid bootstrapping algorithm (-f a) to perform 10,000 bootstrap replicates and ML searches at once (-# 10,000; -T 20). For our Bayesian approach, all prior parameters of MrBayes except for the model settings were kept at default. We started MrBayes with 1,000,000 generations (ngen=1,000,000 samplefreq=5,000; printfreq=5,000; diagnfreq=1,000). Afterwards, parameter values like effective sample size (ESS) were summarized and checked for reliability with Tracer v. 1.7.1 (Rambaut et al., 2018). Trees were summarized with a burn-in of 10%. All phylogenies and their support values were read and plotted with FigTree. Illustrations and modifications were conducted with InkScape. In all analyses, *P. subterraneus* specimens were used as the outgroup.

### Divergence time analysis

For divergence time analysis, we combined the data of 25 *Poecilochirus* specimens with additional Mesostigmata taxa (including our own *Macrocheles* sequences). *Poecilochirus* samples were chosen by the availability and quality of COI and LSU sequence and covered 12 of the genetic clusters. The complete dataset consists of 40 individuals, of which 26 represent the hyporder Parasitiae (one family), 13 the hyporder Dermanyssiae (10 families) and one the infraorder Uropodina (two families), which serves as the outgroup (Supplement Table S2). The analysis was conducted with Beast v. 2.6.3 (Bouckaert et al., 2019). Beast2 includes the Fossilized-Birth-Death Process model (FBD) which is an extended version of the Birth-Death Process (Stadler, 2010; Stadler et al., 2018). Both models assume that every living lineage can experience speciation at rate λ or go extinct at rate μ, but the FBD model allows the treatment of known fossil calibration points as part of the tree prior at node times. Such priors can be set with the graphical user interface Beauti2. We ran Beast2 under the FBD model using fossil data available for five taxa (Supplement Table S3). Monophyly was fixed for samples of the families Parasitidae, Macrochelidae and Digamasellidae, as well as for the infraorder Gamasina, and the superfamilies Dermanyssoidae and Eviphidoidae. Our analysis is based on the COI and LSU genes of which each represents a separate partition. We set the substitution model to be unlinked and determined the GTR and TN93 model for the COI and LSU partition, respectively. The Clock and Tree model were set to be linked and the analysis ran with the Relaxed Clock Log Normal model. We set the five fossil calibration points to the clade nodes where the fossils are assumed to belong, and ran Beast2 with 1,000,000 generations. Trees were stored every 100 generations. Stationarity was reached when all ESS values were above 200 and data were equally distributed in Tracer v. 1.7.1. The final divergence time phylogeny was assembled with TreeAnnotator v2.6.3 (included in Beast2 package) using a burn-in of 10% and the maximum clade credibility as target tree type. Results were plotted using FigTree and edited with InkScape.

### Biogeography and ancestral-area estimation

The divergence time reconstruction was the basis for a biogeographical analysis. We used the R package BioGeoBears (Matzke, 2014) which performs inferences of biogeographic histories on phylogenies. With BioGeoBears, different models of how biogeography may evolve on phylogenies can be tested on a given dated tree. Currently the package includes the Dispersal-Extinction-Cladogenesis model (DEC), a likelihood version of the Dispersal-Vicariance model (DIVALIKE) and the Bayesian Analysis of Biogeography (BAYAREALIKE). Moreover, it provides an extended version for the models by the consideration of additional free parameters like ‘j’ (“jump dispersal”) or ‘x’ (geographical distances) while modeling. The “jump dispersal” parameter simulates the founder-event speciation. It describes that at the time of cladogenesis one daughter lineage inherits the ancestral range, and the other lineage occupies a new area through a rare, long-distance colonization event and founds an instantly genetically isolated population (Matzke, 2014; Zhang et al., 2017). Since the biogeography of *Poecilochirus* is our focus, we pruned the dated tree to a subset containing only *Poecilochirus* specimens (excl. *P. subterraneus*) and used it for the BioGeoBears analysis. We divided the Northern Hemisphere in 6 areas: Western North America (W), Eastern North America (N), Europe (E), Northern Asia + Japan (A), Southern Asia (S), and South East Asia ranging to the Solomon Islands (I). In an initial analysis, we tested whether the existence of only the Bering Land Bridge (BLB) or the North Atlantic Land Bridge (NALB) and the BLB might fit the data better. Three model types were tested in three different versions for each scenario (M0=DEC, DIVALIKE, BAYAREALIKE; M1=DEC+J, DIVALIKE+J, BAYAREA+J, and M2=DEC+J+X, DIVALIKE+J+X, BAYAREA+J+X), and the likelihood and Akaike Information Criterion with sample correction (AICc) were compared between both scenarios. We continued with the scenario showing the highest likelihood (lowest negative log likelihood value: -lnL) and lowest AICc values in most of the models and compared nested models using a Likelihood Ratio Test (same model type: M0 vs. M1 and M1 vs. M2). The AICc was used to compare among model types (DEC vs. DIVALIKE).

A more likely scenario is obtained by running the biogeographical models under a time-stratified analysis. In such an analysis, BioGeoBears can take into account geographical changes and different difficulty levels for dispersal occurring over time. Our time-stratified analysis included three time slices. We tried to represent the geographic conditions of the Eocene/Oligocene, the Miocene, and present conditions. For this scenario we ran the models DEC/DEC+J and DIVALIKE/DIVALIKE+J.

## Results

### Sequence data

We obtained 429 DNA sequences after editing and quality-checking. Of these, 193 COI, 136 ITS, and 79 LSU sequences belonged to mites that resemble *P. carabi*, 10 COI, 6 ITS, and 3 LSU sequences belonged to *P. subterraneus*, and one COI and one LSU sequence were generated from two *Macrocheles* specimens (Supplement Table S1). These sequence data have been submitted to GenBank under the accession numbers MW890765 – MW890966 (COI), MW893012 – MW893060 and MW893063 – MW893153 (ITS), and MW893154 – MW893193 and MW893196 – MW893239 (LSU). The average length of the COI, ITS and LSU sequences are 655bp, 509bp and 645bp, respectively. All sequences that were blasted against the NCBI database matched published sequences from either *Poecilochirus* or Parasitidae, with low E-values and a sequence coverage > 70%. *Macrocheles* samples matched with sequences from *M. nataliae*.

### Identification of genetic clusters

We identified 24 different genetic clusters by the IQtree approach that was based on a concatenated supermatrix of the COI, ITS, and LSU genes obtained from 230 *Poecilochirus* mites (Table 1, Supplement Figure S2). Of these, 3 were clusters in the outgroup *P. subterraneus*. The largest cluster in the ingroup consisted of 89 samples (Europe-1), and seven clusters were represented by only one mite individual (= singletons). Depending on the cluster, the number of different host species ranged from 1 to 8, and the number of sampling locations varied from 1 to 13 (Table 1).

**Table 1:**
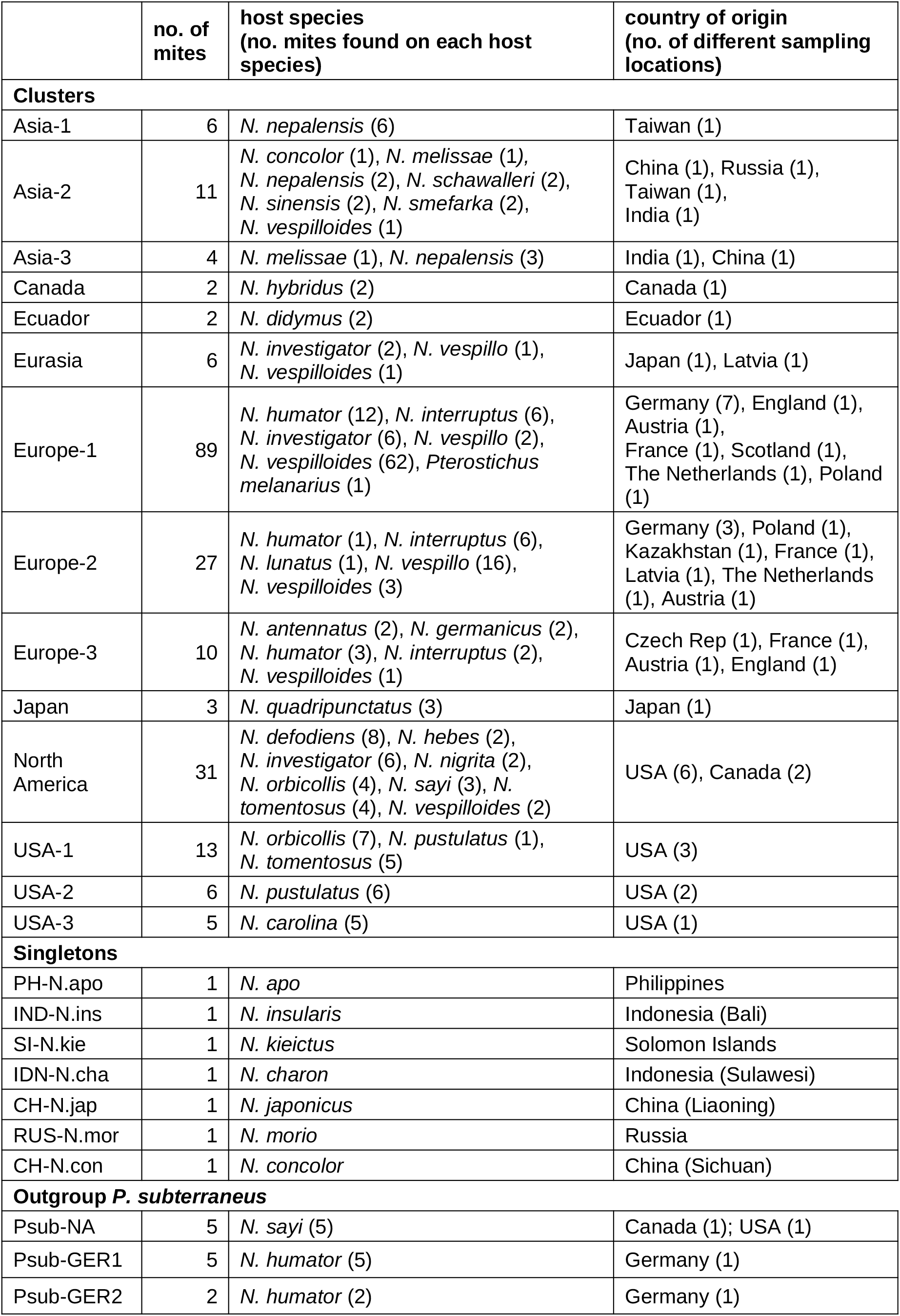
List of genetic clusters of *Poecilochirus*. The number of mite individuals, the host species with the number of mite individuals sequenced from each host species, and the country of occurrence with the number of its different localities are listed for each cluster.

We named the clusters according to their main distribution area. All clusters but Asia-2 were represented by reliable branch support values (SH-aLRT > 80%; UFBoot > 95%). The separation of Asia-1, IDN-N.ins and Asia-2 from the remaining singletons resulted in less confident values (SH-aLRT=26%; UFBoot=53%). However, the close relationships among Asia-1, Asia-2, IDN-N.ins, PH-N.apo and SI-N.kie were well supported as a monophyletic group (SH-aLRT=81.6%; UFBoot=95%). Its sister group is a clade formed by the North America and Eurasia clusters but with lower branch support values (SH-aLRT=71.8%; UFBoot=91%). Branch support values of deeper phylogenetic splits varied highly (SH-aLRT: 20.7--99.9%; UFBoot: 67-100%) indicating a more fragile tree topology (Supplement Figure S2).

The mPTP species delimitation results were similar to those of the IQtree approach. The four independent MCMC runs yielded the highest frequencies for species numbers between 22 and 26 with the highest likelihood score for a multi-coalescent rate (-lnL=1099.35) calculated for 23 species (including 3 *P. subterraneus* clusters, Supplement Figure S3).

Mites in the three *P. subterraneus* clusters were collected in Europe (2 clusters) or North America (Supplement Table S1). The remaining mite clusters were each restricted to one of three major geographical regions (the European, Asian, and American continent). In North America (USA and Canada), five different clusters occurred. While the North America cluster was distributed from Alaska/Canada over the Western to the Eastern USA, the USA-1 cluster was only found in the North-Eastern part of the USA (Illinois, Ohio and Connecticut). The USA-2 cluster was only present in Illinois and Ohio (Middle-Eastern USA), and the USA-3 cluster was restricted to Florida (South-Eastern USA). The Canada cluster only appeared in Calgary/Alberta. In South America the Ecuador cluster was found (Figure 1, Supplement Table S1).

**Figure 1:**
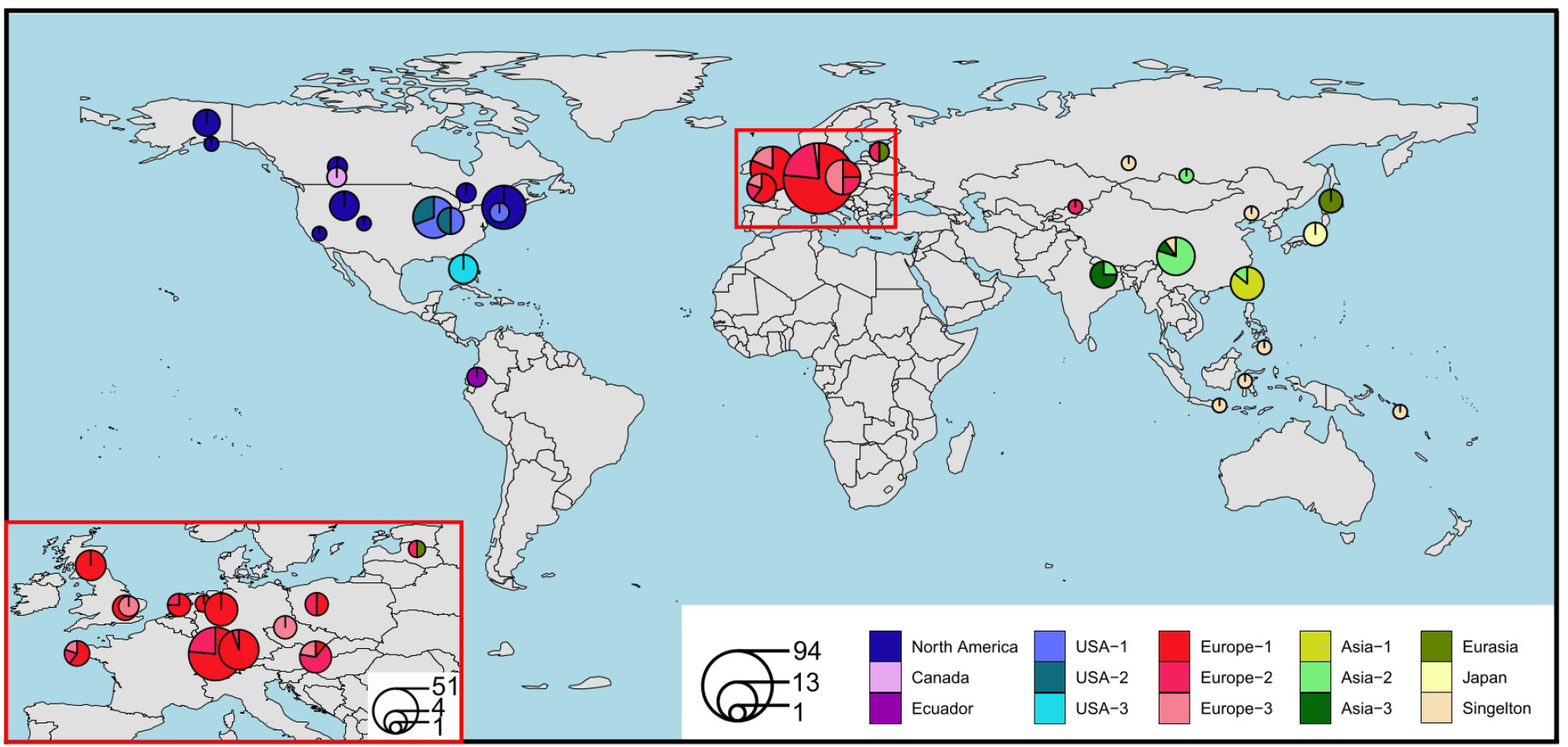
Distribution of the Poecilochirus clusters without P. subterraneus. Pie charts show the relative proportion of the different clusters at each location. Each cluster is represented by another color (except singletons) and pie size reflects the sample size at each location. The European distribution is enlarged in the left bottom corner (red rectangle).

The clusters Europe-1, Europe-2, and Europe-3 were distributed across Europe whereby the Europe-2 cluster also contained a sample from Kazakhstan. Samples of the Eurasia cluster occurred in Latvia and Japan. All singletons and three additional genetic clusters (Asia-1, Asia-2 and Asia-3) were distributed across the Asian continent. We identified a distinct Japan cluster in addition to the Eurasia cluster on the Japanese Islands (Figure 1).

The mean uncorrected p-distance within clusters was 0.78% ranging from 0.1% (Asia-1) to 1.9% (Asia-3). Among clusters the overall mean p-distance was 15.48% with a range between 6.03% (Asia-1/IND-N.ins) and 21.06% (Psub-GER1/USA-2). The p-distance between the known species *P. carabi s*.*s*. and *P. necrophori* (Europe-1 vs. Europe-2) was 10.21%, and that between *P. carabi*/*P. necrophori* and *P. monospinosus* (Europe1/Europe2 vs. USA-2/USA-3) was on average 19.46%.

### Morphological identification

We morphologically identified 95 large *Poecilochirus* specimens covering 19 different genetic clusters. Of these, 90 specimens from 16 genetic clusters match the *P. carabi* description of Hyatt (1980). The specimens from the Japan cluster differed slightly from Hyatt’s description by having a weakly sclerotized body and long podosomal and opisthosomal shields (Supplement Tables S1 and S4).

Five specimens were identified as other species: The single intact specimen of the USA-2 cluster (sample ID: oh-pus2) corresponded to the description of *Poecilochirus monospinosus* Wise, Hennessey & Axtell, 1988. All individuals of the USA-3 cluster resembled *P. monospinosus* as well but differ in the setal pattern. The CH-N.con singleton morphologically resembled *Poecilochirus austroasiaticus* Vitzhum 1930, but it was larger than reported by Hyatt (1980). These morphological results prompted our definition of *P. carabi s*.*l*., which hereafter includes all genetic clusters, except USA-2, USA-3, CH-N.con, and the three *P. subterraneus* clusters.

### Host specificity

We focused on six clusters that contained more than six mite specimens each and that were found in more than one location. Among the three European clusters, Europe-1 and Europe-3 were each associated with five, and Europe-2 with four *Nicrophorus* species, respectively (Table 2; Figure 2).

**Table 2:**
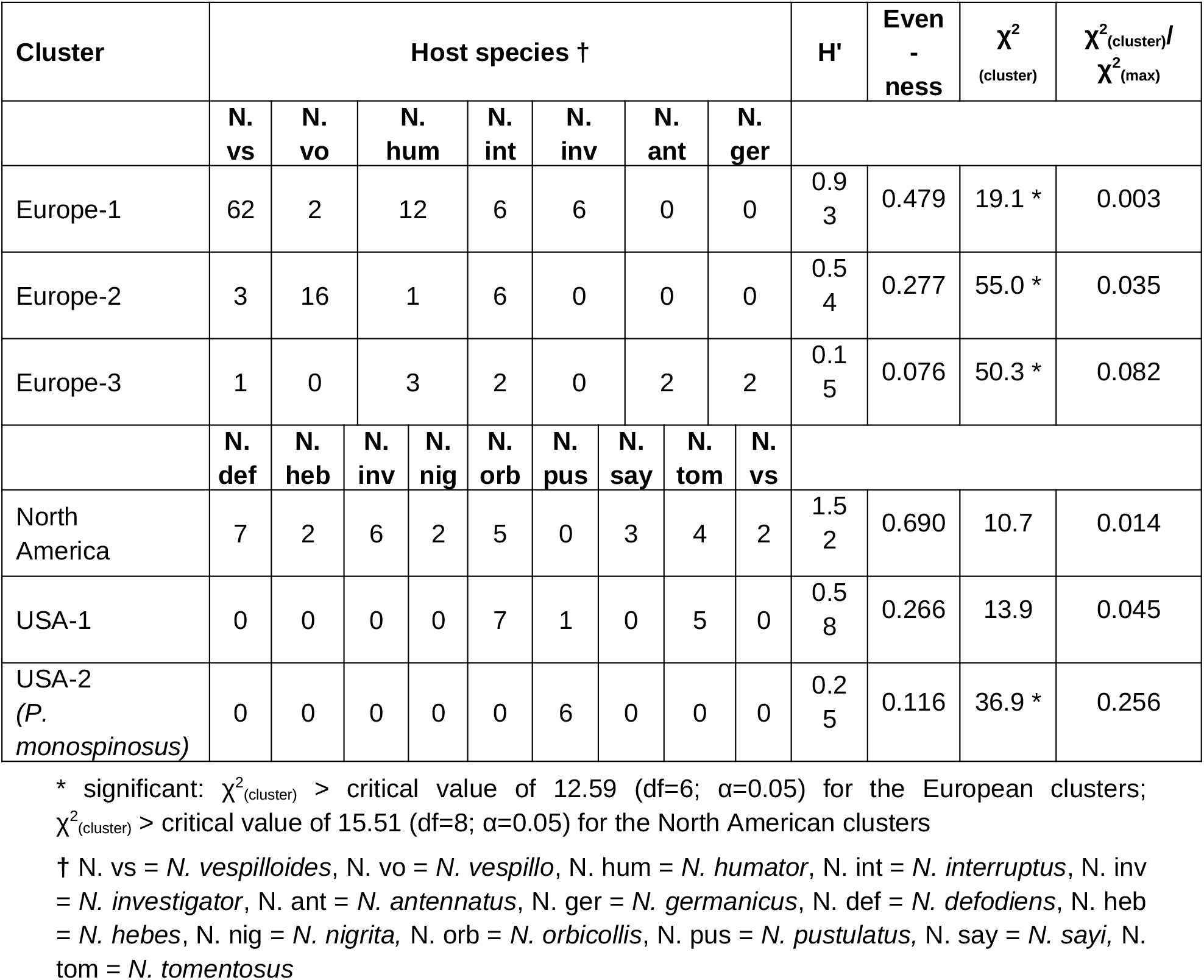
Host specificity indices for three European (n = 124) and three North American (n = 50) clusters. The number of mite specimens found on each host, the Shannon Wiener Diversity Index (H’), Evenness, χ^2^-value and χ^2^-Ratio are listed for each cluster.

**Figure 2:**
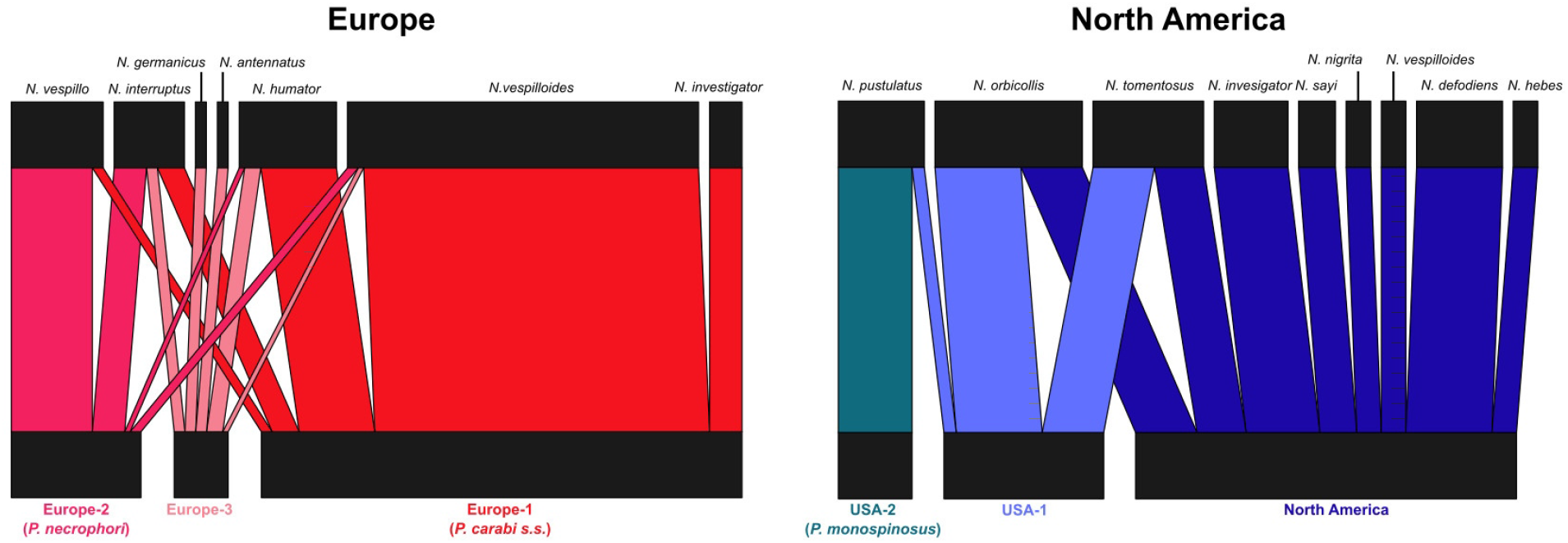
Association network between host species and the six genetic clusters tested for host specificity. The map illustrates the weighted association between mite clusters and *Nicrophorus* species for the clusters Europe-1, Europe-2 and Europe-3, as well as the clusters USA-1, USA-2 and North America. The thicker the bars the more mite individuals are associated with the respective host species.

Both Shannon-Wiener Diversity Index and Evenness were highest for the cluster Europe-1 and lowest for Europe-3 (Table 2). The association between the three North American clusters and their host species is clearer, even though more host species were involved. The clusters USA-2, USA-1 and North America were found on one, three and eight *Nicrophorus* species, respectively (Table 2; Figure 2). Furthermore, a similar pattern for both statistical indices was shown in North America where both values decreased from the North America cluster over USA-1 to USA-2. Furthermore, the χ^2^ ratio ranged from 0.003 to 0.082 in Europe and from 0.014 to 0.256 in North America (Table 2). The higher the quotient, the more the samples from a cluster were concentrated on specific host species.

### Phylogenetic inference

The tree topologies inferred by the ML (IQtree and RaxML) and BI (MrBayes) analyses were consistent. The phylogeny comprised 37 mite individuals covering 16 genetic clusters. Two *P. subterraneus* samples served as the outgroup (Figure 3).

**Figure 3:**
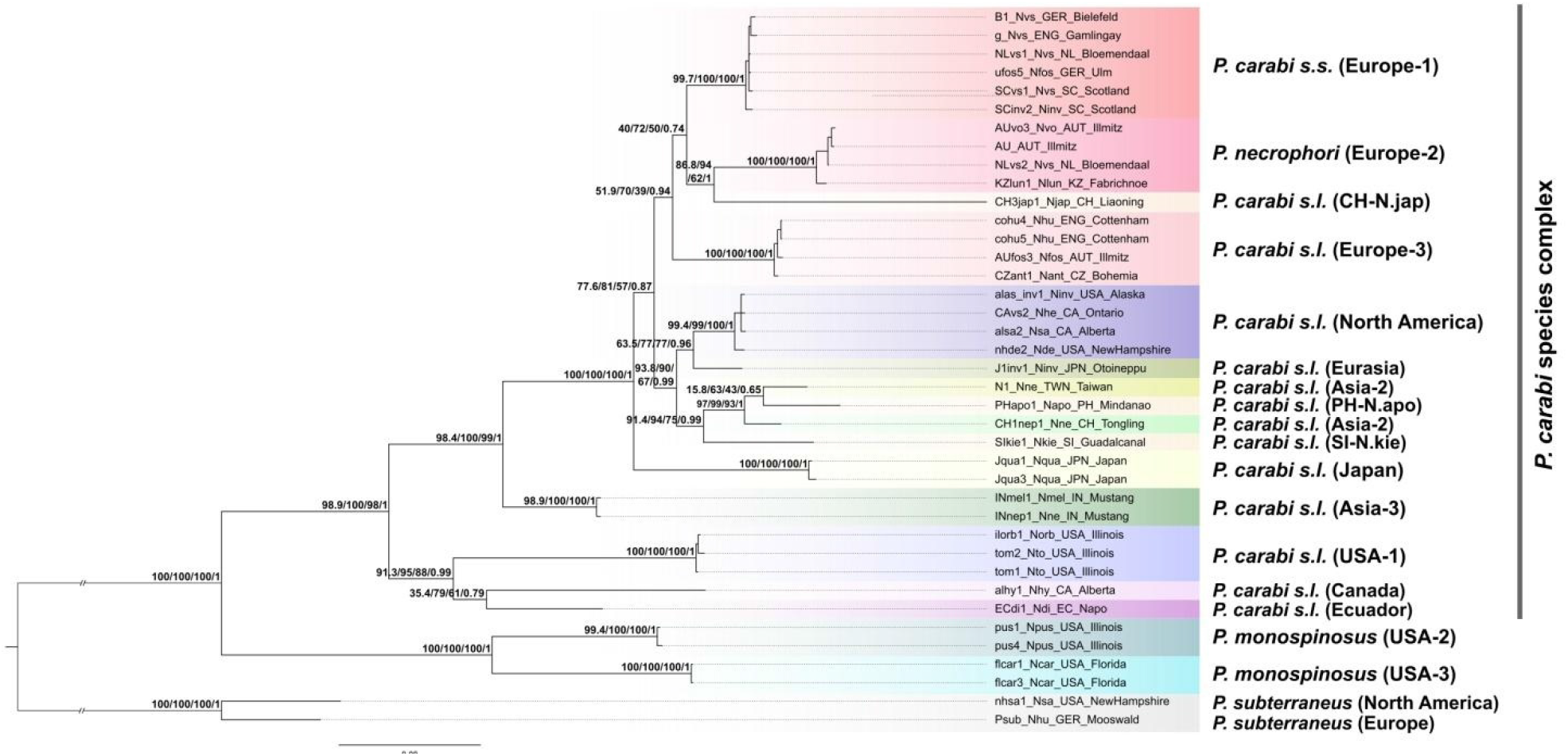
The phylogeny of *Poecilochirus carabi s*.*l*. inferred by MrBayes. Branch labels represent the branch support values obtained by the Likelihood Ratio Test/Ultrafast Bootstrapping/Standard Bootstrapping/Posterior Probability. Genetic clusters are indicated by colors. Basal branches are trimmed and the scale indicates the estimated substitutions per site. Species names are those of the best-fitting species description.

Monophyly of the previously defined genetic clusters was confirmed by SH-aLRT, UFBoot; SBS, and PP values. The topology depicted a basal separation into two clades (PP=1; aLRT/UFBoot/SBS=100). One clade consisted of mites identified likely as *P. monospinosus* (USA-2 and USA-3), while the other one included all clusters of *P. carabi s*.*l*.. Within this *P. carabi s*.*l*. clade, the most recent common ancestor of the USA-1, Canada, and Ecuador clusters split off first but the close relationship between the Canada and Ecuador cluster showed lower branch support (aLRT=35.4; UFBoot=79; SBS=61; PP=0.79). Subsequently, our tree shows first the Asia-3 and then the Japan cluster branching off, both with high support values (PP=1; aLRT/UFBoot/SBS>98). The remaining clusters diverged into two clades but this separation was less confident (aLRT=77.6; UFBoot=81; SBS=57; PP=0.87). While the close relationship between the North America/Eurasia and the Asian cluster had consistent support through all but one branch value (aLRT=93.8; UFBoot=90; SBS=67; PP=0.99), values within the European/CH-N.jap/RUS-N.mor clade were low and varied among analyses (Figure 3).

### Divergence time analysis

The divergence time analysis was based on the COI and LSU gene alignment of 40 specimens covering 10 genetic clusters of *P. carabi s*.*l*., 2 clusters from each *P. monospinosus* and *P. subterraneus*, respectively, and 15 additional Mesostigmata taxa. Certain taxa were represented by chimeric sequences, meaning that the COI and LSU sequences did not originate from the same individual but from the same genus or family (Supplement Table S2).

The phylogenetic tree generated by Beast2 had high support values at all but two branches (split between Phytoseiidae and Podocinidae: PP=0.79, and split among Asia-2 samples: PP=0.81) (Supplement Figure S4). The relaxed-clock model suggested an origin of the Mesostigmata in the Late Jurassic (156.2 Ma; 95% credibility intervals [CI] 77 – 272 Ma; Supplement Figure S4) and the divergence into Parasitiae and Dermanyssiae occurred approximately 40 million years later (114.0 Ma; 95% CI 66 – 189 Ma). Within the Parasitiae clade the first diversification occurred in the early Eocene (53.2 Ma; 95% CI 26 – 90 Ma). The segregation of *P. subterraneus* was suggested to occur in the mid Eocene (43.9 Ma; 95% CI 23 – 74 Ma). The *P. monospinosus* clade branched off during the transition from the Eocene to the Oligocene (34.7 Ma; 95% CI 18 – 59 Ma). Diversification of the *P. carabi s*.*l*. clade started in the Oligocene with the separation of USA-1 (29.5 Ma; 95% CI 15 – 50 Ma). All remaining divergence events occurred during the late Oligocene/Miocene (∼ 5 – 25 Ma; Supplement Figure S4).

### Biogeography and ancestral-area estimation

Regarding the first two scenarios we tested (NALB/BLB vs. BLB-only), the NALB/BLB scenario received higher log-likelihood and lower AICc values than the BLB-only scenario in six out of nine models (Supplement Table S5). Hence, our data better fit the assumption that mites dispersed via both the North Atlantic and Bering Land Bridge. As the BAYAREALIKE model type yielded the lowest percentage value of weighted AICc in both of these scenarios (< 4%; Supplement Table S5), we excluded this model type from further analyses.

Within the NALB/BLB scenario, p-values of the LRT were significant when comparing M0 and M1 (DEC: p=0.04; DIVALIKE: p=0.04), but were non-significant for the M1 and M2 comparison (DEC: p=0.06; DIVALIKE: p=0.08). Hence, the more complex model including the parameter “x” which described the different levels of dispersal difficulties (M2 model) was rejected for both model types.

Regarding the weighted AICc values, DEC+J and DIVALIKE+J yielded the highest percentage with 39% and 45%, respectively (Supplement Table S6). In the third, the time-stratified scenario, the comparison of nested models resulted in an acceptance of the M1 model in all cases (p_(LRT)_<0.05). The AICc and weighted AICc values were lowest and highest, respectively, for the DIVALIKE+J model (Table 3).

**Table 3:** Statistics of the BioGeoBears analysis testing four different models in the time-stratified scenario with log likelihood, likelihood ratio test (LRT; p(LRT)), sample corrected AIC (AICc) and weighted AICc values.

| model | -lnL | LRT | $p_{(LRT)}$ | AICc | # param | # tips | Weighted AICc |
| --- | --- | --- | --- | --- | --- | --- | --- |
| DEC | -35.65 | 4.128 | 0.04 | 75.47 | 2 | 13 | 26% |
| DEC+J | -33.59 |  |  | 75.84 | 3 | 13 | 21% |
| DIVALIKE | -36.5 | 7.191 | 0.001 | 77.17 | 2 | 13 | 11% |
| DIVALIKE+J | -32.91 |  |  | 74.48 | 3 | 13 | 42% |

A dispersal rate of d<0.001, an extinction rate of e=0.55, and a relative per-event weight of founder-event speciation of j=1.28 was estimated. The results suggest that dispersal and vicariance as well as founder-event speciation played an important role for the biogeographic pattern of the mites. The most likely ancestral distribution areas of the time-stratified DIVALIKE+J model are visualized in Figure 4.

**Figure 4:**
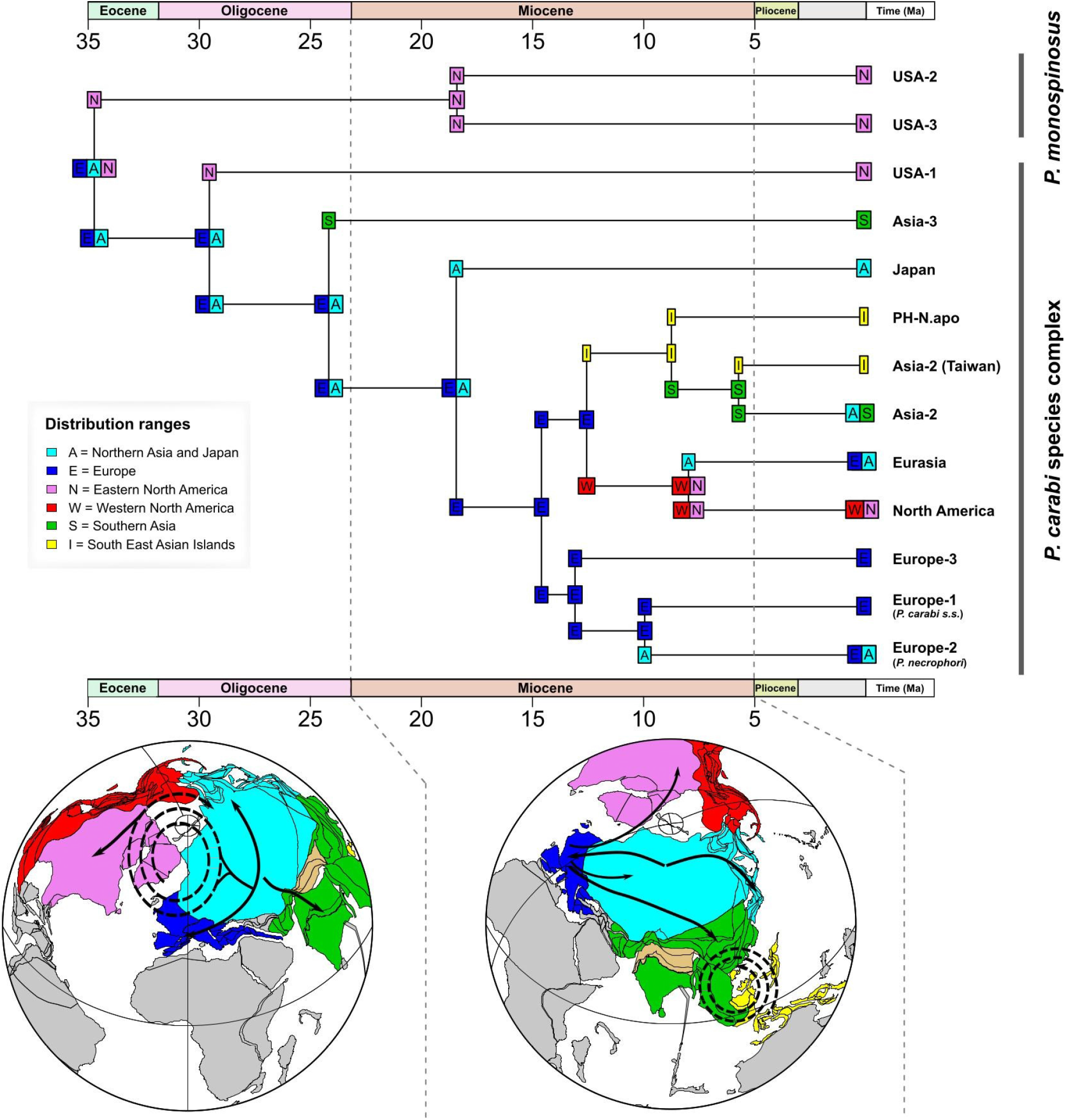
Ancestral state estimation of the DIVALIKE+J model inferred by the time-stratified scenario. Plate tectonic maps are illustrated for 25 Ma and for 12 Ma by the ODSN Plate Tectonic Reconstruction Service (https://www.odsn.de/odsn/services/paleomap/paleomap.html). Black arrows on the maps show dispersals with and without founder-event speciation.

The distribution of the last common ancestor of *P. cabi s*.*l*. and *P. monospinosus* was estimated to range from Eurasia (E and A) to Eastern North America (N). Vicariance was inferred in the branching off of the common ancestor of the USA-2 and the USA-3 cluster (EAN → EA+N), and of the ancestor of the Japan cluster (EA → E+A). Six long-distance dispersals with founder-event speciation were suggested to explain the origin of both the USA-1 and the Asia-3 cluster (EA → EA+N; EA → EA+S), the divergence of the Europe-1/Europe-2 clusters (E → E+A) and the North America/Eurasia clusters (WN → WN+A), and two cladogenesis events within the Asian clade (I → I+S; S → S+I). However, the proportion of the most likely ancestral states deviated just slightly at several cladogenesis events (Supplement Figure S5).

## Discussion

Our study identified 24 distinct genetic *Poecilochirus* clusters including mites that likely represent four different named species: *P. subterraneus, P. monospinosus, P. carabi s*.*s*., and *P. necrophori*. The phylogenetic and species delimitation analyses indicate that many of the genetic clusters found within these morphological species are likely to be cryptic species, most of which are to our knowledge not formally described. We cannot infer with certainty the geographical origin of *Poecilochirus* with our data set, but mites appear to have migrated more than once between Asia, Europe and North America. We also found indication that some mite clusters are specialized on particular *Nicrophorus* species, which may have driven speciation, but this pattern appears to be largely concealed by the effects of multiple migrations between continents. It is difficult to separate these interwoven factors in the evolution of mite species, obfuscating our understanding of the importance of coevolution with hosts and sympatric speciation in *Poecilochirus*. However, we can state with certainty that all speciation events we could infer happened more than five million years ago, with no indication of recent speciation events or ongoing segregation among extant populations.

### Cryptic diversity and host specificity of *Poecilochirus* mites

Cryptic species have been uncovered by molecular investigations across many mite groups (Beaurepaire et al., 2015; Knee, Beaulieu, Skevington, Kelso, & Forbes, 2012; Knee et al., 2012; Schäffer & Koblmüller, 2020) and based on our molecular analyses, we propose that *P. carabi s*.*l*. consists of at least 17 genetic clusters. Genetically, samples within each cluster are very similar (p-distance < 2%), but clusters differ clearly and consistently from each other with a COI divergence of at least 6% between clusters. Given that the well-studied biological species *P. necrophori* and *P. carabi s*.*s*. (Baker & Schwarz, 1997; Müller & Schwarz, 1990; Nehring et al., 2017) are relatively closely related within *P. carabi s*.*l*. and diverge by 10.21% in their COI sequences, we suggest that most of the genetic clusters that we document, at least those for which we sequenced enough replicates (e.g. Europe-3 or North America), represent separate biological species, most of which are undescribed.

### Poecilochirus subterraneus

*Poecilochirus subterraneus* served as the outgroup in our study. The species has previously been observed in Europe (mostly on different *Nicrophorus* species; Hyatt, 1980; Korn, 1983), North America (*N. investigator, N. nigrita*; Grossman & Smith, 2008; Sikes et al., 1996), and Asia (*N. quadripunctatus*, Satou et al., 2000). Here, we sequenced American and European mites resembling the *P. subterraneus* description and found that while mites from both continents clustered together in our dataset, they segregated into three distinct clusters, one from North America and two from Europe. While species delimitation may be unreliable because of limited sampling among the *P. subterraneus* clusters, our data indicate that *P. subterraneus* might be more diverse than previously thought.

### The clades *P. monospinosus* (USA-2 & USA-3) and *P. cf. austroasiaticus* (CH-N.con)

The mites that morphologically resemble the description of *P. monospinosus* fall into two separate genetic clusters that we found on two different host species. USA-3 (n=5) have been sampled from *N. carolina* in Florida only. We did not have any samples available from other beetle species in Florida, thus we cannot speculate about any potential specialization on *N. carolina*.

We found USA-2 mites only associated with *N. pustulatus*. This host association persisted across two locations. In both, *N. pustulatus* occurs sympatrically with other host species (*N. orbicollis, N. tomentosus*). In our data set, only one mite from another genetic cluster was found on *N. pustulatus*. USA-2 was also the cluster with the second lowest Shannon-Wiener index and evenness, and by far the highest χ^2^ ratio, a measure that takes into account the sampling data quality of the cluster in question. USA-2 thus appears to be a strict monospecific host specialist on *N. pustulatus. Nicrophorus pustulatus* has a unique ecology; it has been reported to reproduce on snake eggs and in bird nests on dead nestlings, and it occurs predominantly in the forest canopy, while other *Nicrophorus* species are typically found near the ground (DeMarco & Martin, 2020; Smith et al., 2007; Wettlaufer et al., 2018). *Nicrophorus pustulatus* thus occupies a distinct ecological niche that may isolate the mites, and possibly select for adaptations specific to this niche. Other families of phoretic mites associated with *N. pustulatus* showed no apparent host specificity for this beetle species (Knee, 2017; Knee et al., 2012), indicating that this beetle’s unique niche has not caused mite divergence in every case. Previously, *P. monospinosus* had only been described from poultry manure, preying on fly eggs and larvae – it has not been documented on beetle hosts (Wise et al., 1988). This begs the question whether mites of the original description are an aberrant lineage without beetle association or whether *P. monospinosus* is more general in its host usage and occurs with and without beetles.

The mite individual from *N. concolor* found in Central China (Sichuan Province) is particularly interesting because morphologically it resembles *P. austroasiaticus* more than *P carabi s*.*s*. or any other described species. A discovery of this species in Central China and the association with *N. concolor* is an unexpected observation as so far *P. austroasiaticus* has only been recorded in Siberia/Northwestern China and Europe on animal corpses, or in association with silphid beetles including *N. investigator* (Hyatt, 1980; Makarova, 2013). According to the phylogenetic approach of cluster identification (IQtree analysis), this singleton is closely related to the clade of *P. monospinosus* (Supplement Figure S2).

### The European clusters: *P. carabi s*.*s*., *P. necrophori*, and a new clade

We found three clusters that are almost exclusively distributed in Europe and are closely related. We can unequivocally assign the clusters Europe-1 and Europe-2 to the species *P. carabi s*.*s*. and *P. necrophori* because we tested the host preference of some mites before killing them (Schwarz, 1996). Based on their association with different beetle species, these mite clusters appear to either prefer or to avoid certain hosts, which is in agreement with observations on the host range of the two reproductively isolated mite species (Müller & Schwarz, 1990; Nehring et al., 2017; Schwarz, 1996; Schwarz et al., 1998). Several *Nicrophorus* species occur sympatrically in Europe, and most of them overlap in their seasonal reproductive period but differ in seasonal and diel activity and habitat use (Dekeirsschieter et al., 2011; Esh & Oxbrough, 2021; Majka, 2011; Müller & Eggert, 1987; Schwarz & Koulianos, 1998; Scott, 1998). Thus, as earlier work already suggested (Schwarz, 1996), the European mite generalist (Europe-1 = *P. carabi s*.*s*.) exploits several host species with different life history traits, while *P. necrophori* (= Europe-2) is specialized on *N. vespillo*, which prefers open area habitats where other *Nicrophorus* species are less abundant (Esh & Oxbrough, 2021).

We found a third cluster of mites from across Europe: Europe-3. At the moment, this cluster is a curious case because it is widespread across Europe but was not found in Germany, where most of our samples were collected. The three European clusters were found in sympatry in some locations (France, Austria), indicating that they do not necessarily competitively exclude each other. Sun and Kilner (2019) described a *P. carabi s*.*l*. population from the United Kingdom that differs in its phenotype from *P. carabi s*.*s*.. While this population may correspond to *P. necrophori*, it is tempting to speculate that it is in fact our Europe-3 cluster, given that we did not find any *P. necrophori* among the 16 mites from the UK.

### American samples of *P. carabi s.l*

The ecology and behavior of the North American *P. carabi s.l*. is quite well studied, although not to the same extent as the European populations. Brown & Wilson (1992) reported two reproductively isolated populations from Michigan that differed in morphology, and their preference for host species of *Nicrophorus*. We were not able to obtain any reference samples from Michigan but our analysis confirms the occurrence of at least two genetic clusters from Northeastern America (USA-1, North America) in addition to *P. monospinosus*. We also found evidence for further clusters from Canada and South America.

The North America cluster was the most diverse in terms of host species numbers, and the cluster did not appear to prefer any specific host species among those occurring across its distribution range. Such a broad host range increases the independence of host abundance, seasonal and diel activity, and other life history traits. In comparison, USA-1 was found almost exclusively on *N. orbicollis* and *N. tomentosus* in the northeastern USA, which might be an indication of local specialization on two host species. Being a local specialist on two sympatric *Nicrophorus* beetles with different seasonal activity (Brown & Wilson, 1992; Keller et al., 2019; Scott, 1998; Wilson, 1982) could expand the reproductive period of the mites.

Some previously described populations of *P. carabi s.l*. from Michigan indeed reproduced successfully using various host species, but others were local specialists (Brown & Wilson, 1992, 1994; Wilson, 1982). In our data, we found no evidence of any clusters being strict specialists for either *N. orbicollis* or *N. tomentosus*, as had been reported by Brown and Wilson (1992) for the Michigan populations. While it is possible that these Michigan populations belong to genetic clusters that we did not sample, it may also be that genetic clusters are specialists in one community and less strict in their host choice in another (Brown & Wilson, 1992).

### Asian diversity

Our Asian samples cover a great number of sampling locations and host species, but we could only analyze a few or one replicate individuals for most of the genetic clusters. Thus, any ecological inference is impossible and we may have only scratched the surface of the biodiversity of *Poecilochirus* mites that use *Nicrophorus* as hosts in Asia.

### Phylogenetic inference

Our phylogeny provides a basic overview of the relationships among *Poecilochirus* mites. Maximum likelihood and Bayesian Inference analyses reveal congruent tree topologies without polytomous relationships among the clusters. We applied different branch support methods (SH-aLRT, UFBoot, SBS, PP) as their accuracy is debated and confidence levels can vary (Anisimova et al., 2011; Pyron et al., 2011). Relationships between clusters are resolved at deeper levels and in some derived clades, as indicated by well-supported branches across all methods. Support values of a medium range (50 < SH-aLRT/UFBoot/SBS < 75; 0.5 < PP < 0.75) occur mainly in more derived relationships and reveal higher values for posterior probabilities than for bootstrap approximations. Such deviations occur because bootstrap values are a more conservative support measure than Bayesian posterior probabilities which can produce a higher false-positive rate (Anisimova et al., 2011; Cummings et al., 2003; Erixon et al., 2003). According to the variation of support values at some branches, certain phylogenetic relationships between European, Asian, and North American clusters should be interpreted with caution. Low support values could result from inconsistencies between the gene trees as we used a concatenated supermatrix of COI, ITS, and LSU, with a partitioning approach. Branch lengths are analogous across the analyses and express an adequate amount of genetic change between internal nodes. Certain branches are in fact relatively long (e.g., USA-2, USA-3 and CH-N.jap). This high level of genetic change could be due to an accelerated rate of evolution, a high extinction rate leaving only one member of a radiation, or an underrepresented diversity caused by incomplete taxon sampling.

However, the congruent tree topology inferred by all analyses, the medium to high support values, the appropriate branch lengths, and the exclusive dichotomy all indicate a high degree of robustness for this phylogeny. The genes we concatenated for this phylogenetic reconstruction already provided sufficient genetic information individually to distinguish between morphologically described species in other groups of mites (Lehmitz & Decker, 2017; Lv et al., 2014; Schäffer & Koblmüller, 2020). In general, the mites showed low morphological variability, despite their high genetic divergence, but the combined approach of molecular and morphological techniques helps us to better understand the species boundaries and cryptic diversity in this unique group of mites.

### Evolutionary history and biogeography

Our data suggest a split between the ancestors of *P. carabi s.l*. and the *P. monospinosus* clade during the Eocene/Oligocene and a further radiation within *P. carabi s.l*. in the Miocene. During this period, most of the *Nicrophorus* diversity already existed (Sikes & Vernables, 2013). Although the geographic origin of their common ancestor cannot be stated with certainty, the ancestral area of the *P. monospinosus* clade is clearly the North American continent. *Poecilochirus carabi s.l*. might have originated in Eurasia with an early dispersal to the New World (USA-1). The likelihood proportions of the most likely ancestral areas differ only slightly at this cladogenesis event (Supplementary Figure S5), but regardless of ancestral area, the mites moved between the New and Old World during the Eocene/Oligocene. The Miocene diversification of *P. carabi s.l*. took place in Eurasia with at least one colonization of the New World that is less debatable in terms of dispersal direction (resulting in the North America clade). In both epochs (Eocene/Oligocene and Miocene), a connection between Eurasia and North America by the Bering and North Atlantic land bridges is assumed (Brikiatis, 2014; Denk et al., 2010; Graham, 2018; Jiang et al., 2019; Tiffney, 1985). Although the Bering Land Bridge is often considered the only relevant connection between the continents for floral and faunal migration (Lee et al., 2020; Wen et al., 2016), the assumption that the mites used both land bridges fits our data better. Hence, a closer look at the phylogenetic relationships of beetles and mites occurring near the North Atlantic (e.g., Western Europe; Eastern Canada) would be useful in assessing the role of a North Atlantic Land Bridge and its suitability for the dispersal of small organisms. Regardless of the routes on which the mites migrated between continents, Europe might be a pivotal starting point for their dispersal during the Miocene.

In Southern Asia, mites colonized multiple areas. As this region experienced several geological and climatic changes since the early Miocene that could have resulted in the origin of new geographical barriers (e.g., sea level changes and aridification: Bird et al., 2005; Miao et al., 2012; Zhisheng et al., 2001), vicariant speciation might have contributed to the scattered pattern of Asian clusters.

We would like to emphasize that models including the “jump dispersal” parameter were most-likely in all biogeographic scenarios, which highlights the importance of founder-event speciation for the evolution of *Poecilochirus* mites. Furthermore, the results of the tested biogeographic model types DEC+J and DIVALIKE+J deviate just slightly among all analyses. This indicates that dispersal with extinction and vicariance are key processes for understanding the historical biogeography of this species complex.

### Drivers of speciation

Based on our phylogenetic analyses, geographic separation is responsible for the main divergence among *P. carabi s.l*. lineages, while the specialization on certain beetle species did not play a significant role on a global scale. Although host-parasite relationships are seen as an important driver of sympatric speciation, co-speciation with the host beetles can be largely ruled out for the *Poecilochirus* in our study since the main mite diversification happened at least 40 million years after the radiation of the burying beetles (∼ 75 Ma, Sikes & Venables, 2013).

Because the mites are small and cannot fly, the widely separated distribution areas of the mite clusters are likely due to beetle mobility rather than mite mobility. This may have also facilitated founder events since beetles typically carry several mites, thus increasing the probability that individuals arriving in new areas will be able to find a mate. The holarctic species *N. vespilloides* and *N. investigator* are of particular interest as both species dispersed either from the New to the Old World (*N. vespilloides*) or vice versa (*N. investigator*) (Sikes et al., 2008; Sikes & Venables, 2013). Mites carried by these beetle species appear in multiple genetic clusters, some of which are also closely related (e.g. Eurasia and North America cluster). This suggests that both *Nicrophorus* species played a major part in the dispersal and evolution of *P. carabi s.l*..

Within the continents, further radiation seems to have taken place, with some lineages specializing on certain hosts (e.g USA-2, Europe-2) and others having a broader host range (Europe-1, North America). In this context, European populations may have radiated in sympatry. Their hosts use different microhabitats to which the mites may have adapted. For example, *N. vespillo*, host to the specialized *P. necrophori*, is more common in meadows, while *N. vespilloides* is more abundant in forested areas in Germany, the UK, and Alaska (Majka, 2011; Scott, 1998, Sikes et al., 2016). This may have caused genetic divergence among the mites, both through drift after spatial separation and ecological adaptation. Since meadows are more sun-exposed, *N. vespillo* and its mites may be adapted to warmer temperatures. When kept at the same temperature, *N. vespilloides* develops quicker than *N. vespillo*, which may be the result of countergradient variation across the two species (Conover & Schultz, 1995; Müller & Schwarz, 1990). The mites they carry track this difference in their own development time, which may either be a direct adaptation to temperature or an indirect one - because mite development needs to be completed before beetle development for optimal dispersal (Müller & Schwarz, 1990; Nehring et al 2017; see Brown & Wilson, 1992 for a similar effect in American populations). Selection on mite development time (e.g. in the event of a host switch) can lead to rapid adaptation in development time and correlated changes in other traits through hitchhiking or pleiotropy (Schedwill et al., 2018). These effects could cause reproductive isolation among the differentially selected mite populations (Nosil & Harmon, 2009), and thus host specialization can drive genetic divergence. Indeed, a relationship between genetic divergence and host specificity has also been reported from other parasites, such as the honey bee parasite *Varroa* (Beaurepaire et al., 2015) and *Macrocheles* species that are associated with *Nicrophorus* beetles (Knee, 2017). Genetic clustering driven by host adaptation is also found in feather mites that are phoretic on seabirds (Stefan et al., 2018). Host specificity is often seen as a species-level trait. However, it should be considered that local populations of one and the same species could encounter different host communities and may thus specialize on different hosts, an excellent subject for future studies (Brown & Wilson, 1992; Korallo-Vinarskaya et al., 2009; Thompson, 2009).

## Conclusions

Our global analysis of the *P. carabi* species complex revealed a surprisingly high genetic diversity. Our data support previous ecologically and morphologically defined species clades for *P. necrophori, P. carabi s.s*., *P. monospinosus*, and *P. cf. austroasiaticus*. In general, phylogenetic relationships between the mite clusters did not match those of the beetles (Sikes & Venables, 2013).

Drivers of genetic diversification differ depending on the geographic scale. Globally, spatial separation between continents can explain the deep splits between clades relatively well, although back-migrations among continents are obvious. Within continents, further diversification appears to have occurred that was independent of separation by oceans (e.g. among Europe 1-3). While this may have happened due to spatial separation on smaller scales that we cannot track, it may have also been driven by ecological factors like adaptation to different host species. Separate mite clusters may evolve in sympatry by ecological adaptation directly to local hosts or to the abiotic environment the hosts live in. The former is most likely for the apparent host specialists that we have identified in our dataset. It is still uncertain in which cases the mites may adapt to the host species or just take any opportunity for dispersal and/or reproduction. However, the close association and occurrence with their beetle hosts shaped the evolution of *Poecilochirus* mites, and putative drivers depend on the host communities as well as the features of their biotic and abiotic environment.A taxonomic revision of the genus in future investigations would greatly facilitate our understanding of speciation and biogeographic processes in this species complex and genus.

## Acknowledgments

We are indebted to all colleagues who sent us mites they had collected: A Barsevskis, M Brandley, J&I Brookhart, JM Carpenter, L DeCicco, R Enser, J Hajek, B Jarrett, L Jingke, Z Kaszab, R Kilner, G Kushnak, T Larsen, R Leech, J Longino, R Madge, K Maruyama, M Maruyama, D Mohagan, T Mousseau, JK Müller, P Naskrecki, M Nishikawa, S Peck, J Peck, C Raithel, A Riedel, J Ruzicka, K Sagata, M Schilthuizen, MP Scott, KS Sheldon, H Sipkova, J Smiley, PT Smiseth, RJ Smith, S Steiger, H Sugaya, S Suzuki, M Ulyshen, A Urbański, D Wagner, S Werner, J Withrow, N Wood. For help with expeditions to China, Nepal, and Japan we graciously thank: P Naskrecki, and R Ziedler (China), D Manandhar, and S Peck (Nepal) and S Suzuki, M Ôhara, T Nisimura, M Maruyama, and M Nagano (Japan). Funding to VN was provided by the Freiburg Research Innovation Fund and the German Research Foundation (NE 1969/3-1), and the Rapid Assessment Program (RAP) of Conservation International, a NSERC Discovery grant, a National Science Foundation Grant (DEB-9981381), a University of Connecticut Research Council grant, and a National Geographic Society grant supported DSS’s field work. We thank Karen Meusemann and Josef K Müller for insightful discussions.

## Supplemental information

**Supplement Table S1:** Overview of single samples with cluster assignment, sampling time and location, host species, country, storage, morphological affiliation, and cluster affiliation.

**Supplement Table S2:** Downloaded COI and LSU sequences of the Mesostigmata taxa with accession number and references. Sequences are used for the divergence time analysis.

**Supplement Table S3:** Fossil data used in the divergence time analysis. Listed are the taxonomic state (superfamily, family and/or genus), age in million years ago [Ma].

**Supplement Table S4:** Morphometric measurements of specimens underwent DNA extraction. Measurements are listed for the podosomal length and width, sternal plate length, opithosomal length, the z1, s1, s2, r3, st4, the macroseta leg IV, and the length of digitus mobilis. All measurements are in µm.

**Supplement Table S5:** Comparison table of likelihood and sample corrected Akaike Information Criteria values (AICc) for the NALB/BLB and BLBonly scenarios inferred by the BioGeoBears analysis.

**Supplement Table S6:** Statistics of the BioGeoBear analysis to test for the best fitting biogeographic model considering the scenario NALB/BLB. Log likelihood values for each model are calculated and nested models (M0 vs. M1 and M1 vs. M2) are tested using the Likelihood ratio test (LRT) with its p-values. Non-nested model comparisons are evaluated by the sample corrected Akaike Information Criteria (AICc).

**Supplement Figure S1:** Sample distribution. Coloration of the world map indicates the annual temperature obtained from WorldClim data (Bioclimatic variable BIO1 – Annual Mean Temperature, www.worldclim.org). Green shaded areas represent the current distribution area of *Nicrophorus* and each dot represents one sample locality of the mite sample set.

**Supplement Figure S2:** Phylogenetic reconstruction of the entire *Poecilochirus* dataset obtained with IQtree. Branch labels represent SH-aLRT and ultrafast bootstrap values in percentage. High branch support is suggested with SH-aLRT > = 80% and UFBoot > = 95%. Clades which show short branch length and/or polytomy have been collapsed. Clade/Cluster labels are assigned to the right.

**Supplement Figure S3:** Species delimitation analysis performed with the mPTP tool. The analysis is based on the entire *Poecilochirus* data set and its resulting IQtree phylogeny. The analysis inferred a total of 23 species (including the 3 *P. subterraneus* species). The diagram shows the likelihood distribution of the 4 MCMC runs over 10 million iterations.

**Supplement Figure S4:** Time-calibrated tree of the Mesostigmata taxa with the focus on divergence times of the *P. carabi* species complex generated by Beast2. Represented branch labels indicate posterior probabilities below 0.9, branches without labeling have support values above 0.9. The chronogram tree shows mean node ages and node bars represent a 95% credibility interval. Time scale on the bottom refers to millions of years ago. Red circles depict fossil calibration points.

**Supplement Figure S5:** Illustration of the ancestral state estimation with proportions of all ancestral states at each node depicted as pie charts. Values are calculated with the R package BioGeoBears and represent the results of the DIVALIKE+J model conducted by the time-stratified analysis. The legend shows all possible ancestral state combinations allowed by the analysis settings.

## Author contributions

JC conducted bioinformatic analyses, conceptualized and drafted the manuscript.

AKE, WK, DSS edited the manuscript

JC, PH, WK, MN, NS conducted molecular wet lab work and initial sequence analyses

JB identified and measured mite specimens

WH, PH, WK, VN, NS, DSS contributed mite specimens

VN conceived and supervised the project, co-wrote the manuscript

## Data Accessibility

DNA sequences: Genbank accessions MW890765 – MW890966 (COI), MW893012 – MW893060 and MW893063 – MW893153 (ITS), and MW893154 – MW893193 and MW893196 – MW893239 (LSU) are provided at NCBI.

## References

Anderson, R. S. 1982. Resource partitioning in the carrion beetle (Coleoptera: Silphidae) fauna of southern Ontario: ecological and evolutionary considerations. Canadian Journal of Zoology 60: 1314–1325.

Anisimova, M., Gil, M., Dufayard, J. F., Dessimoz, C., & Gascuel, O. (2011). Survey of branch support methods demonstrates accuracy, power, and robustness of fast likelihood-based approximation schemes. Systematic Biology, 60(5), 685–699.

Aoki, J. (1974) On the fossil mites in Mizunami amber from Gifu Prefecture, Central Japan. Bulletin of the Mizunami Fossil Museum, 1, 397–399.

Arribas, P., Andújar, C., Moraza, M. L., Linard, B., Emerson, B. C., & Vogler, A. P. (2020). Mitochondrial metagenomics reveals the ancient origin and phylodiversity of soil mites and provides a phylogeny of the Acari. Molecular Biology and Evolution, 37(3), 683–694.

Baker, A. S., & Schwarz, H. H. (1997). Morphological differences between sympatric populations of the Poecilochirus carabi complex (Acari Mesostigmata: Parasitidae) associated with burying beetles (Silphidae: Nicrophorus). Systematic Parasitology, 37, 179–185.

Beaurepaire, A. L., Truong, T. A., Fajardo, A. C., Dinh, T. Q., Cervancia, C., & Moritz, R. F. A. (2015). Host specificity in the honeybee parasitic mite, Varroa spp. in Apis mellifera and Apis cerana. PLoS ONE, 10(8), 1–12.

Bird, M. I., Taylor, D., & Hunt, C. (2005). Palaeoenvironments of insular Southeast Asia during the Last Glacial Period: a savanna corridor in Sundaland? Quaternary Science Review, 24(20–21), 2228–2242.

Bouckaert, R., Vaughan, T. G., Barido-Sottani, J., Duchêne, S., Fourment, M., Gavryushkina, A., Heled, J., Jones, G., Kühnert, D., De Maio, N., Matschiner, M., Mendes, F. K., Müller, N. F., Ogilvie, H. A., Du Plessis, L., Popinga, A., Rambaut, A., Rasmussen, D., Siveroni, I., … Drummond, A. J. (2019). BEAST 2.5: An advanced software platform for Bayesian evolutionary analysis. PLoS Computational Biology, 15(4), 1–28. https://doi.org/10.1371/journal.pcbi.1006650

Brikiatis, L. (2014). The de geer, thulean and beringia routes: Key concepts for understanding early cenozoic biogeography. Journal of Biogeography, 41(6), 1036– 1054.

Brown, J. M. (1989). Specialization in Poecilochirus carabi, a phoretic mite. Michigan State University, East Lansing, Michigan, USA.

Brown, J. M., & Wilson, D. S. (1992). Local specialization of phoretic mites on sympatric carrion beetle hosts. Ecology, 73(2), 463–478.

Brown, J. M., & Wilson, D. S. (1994). Poecilochirus carabi: Behavioral and Life-History Adaptations to Different Hosts and the Consequences of Geographical Shifts in Host Communities. In M. A. Houck (Ed.), Mites: Ecological and Evolutionary Analyses of Life-History Patterns (1st ed., pp. 3–22). Capman & Hall Inc.

Burke, K., Wettlaufer, J., Beresford, D., & Martin, P. (2020). Habitat use of co-occurring burying beetles (genus Nicrophorus) in southeastern Ontario. Canadian Journal of Zoology, 98(9), 591–602.

Butlin, R., Debelle, A., Kerth, C., Snook, R. R., Beukeboom, L. W., Castillo Cajas, R. F., Diao, W., Maan, M. E., Paolucci, S., Weissing, F. J., van de Zande, L., Hoikkala, A., Geuverink, E., Jennings, J., Kankare, M., Knott, K. E., Tyukmaeva, V. I., Zoumadakis, C., Ritchie, M. G., … Schilthuizen, M. (2012). What do we need to know about speciation? Trends in Ecology and Evolution, 27(1), 27–39.

Conover, D. O., & Schultz, E. T. (1995). Phenotypic similarity and the evolutionary significance of countergradient variation. Trends in Ecology and Evolution, 10(6), 248–252.

Cummings, M. P., Handley, S. A., Myers, D. S., Reed, D. L., Rokas, A., & Winka, K. (2003). Comparing Bootstrap and Posterior Probability Values in the Four-Taxon Case. Systematic Biology, 52(4), 477–487. https://doi.org/10.1080/10635150309311

De Gasperin, O., & Kilner, R. M. (2015). Friend or foe: Inter-specific interactions and conflicts of interest within the family. Ecological entomology, 40(6), 787–795.

Dekeirsschieter, J., Verheggen, F., Lognay, G., & Haubruge, E. (2011). Large carrion beetles (Coleoptera, Silphidae) in Western Europe: A review. Biotechnologie, Agronomie, Société et Environnement, 15(3), 435–447.

DeMarco, K. V. B., & Martin, P. R. (2020). A case of a Pustulated Carrion Beetle (Nicrophorus pustulatus, Coleoptera: Silphidae) burying live Tree Swallow (Tachycineta bicolor, Passeriformes: Hirundinidae) nestlings under the nest. The Canadian Field-Naturalist, 134(3), 217–221.

Denk, T., Grímsson, F., & Zetter, R. (2010). Episodic Migration of Oaks to Iceland: Evidence for a North Atlantic “Land Bridge” in the latest Miocene. American Journal of Botany, 97(2), 276–287.

Dunlop, J. A., Kontschán, J., Walter, D. E., & Perrichot, V. (2014). An ant-associated mesostigmatid mite in Baltic amber. Biology Letters, 10(9), 20140531.

Dunlop, J. A., Kontschán, J., & Zwanzig, M. (2013). Fossil mesostigmatid mites (Mesostigmata: Gamasina, Microgyniina, Uropodina), associated with longhorn beetles (Coleoptera: Cerambycidae) in Baltic amber. Naturwissenschaften, 100(4), 337–344.

Esh, M., & Oxbrough, A. (2021). Macrohabitat associations and phenology of carrion beetles (Coleoptera: Silphidae, Leiodidae: Cholevinae). Journal of Insect Conservation, 25(1), 123–136. https://doi.org/10.1007/s10841-020-00278-4

Erixon, P., Svennblad, B., Britton, T., & Oxelman, B. (2003). Reliability of Bayesian Posterior Probabilities and Bootstrap Frequencies in Phylogenetics. Systematic Biology, 52(5), 665–673. https://doi.org/10.1080/10635150390235485

Farish, D. J., & Axtell, R. C. (1971). Phoresy redefined and examined in Macrocheles muscaedomesticae (Acarina: Macrochelidae). Acarologia, 13(1), 16–29.

Folmer, O., Black, M., Hoeh, W., Lutz, R., & Vrijenhoek, R. (1994). DNA primers for amplification of mitochondrial cytochrome c oxidase subunit I from diverse metazoan invertebrates. Molecular Marine Biology and Biotechnologiew, 3(5), 294–299.

García-Varela, M., De León, G. P. P., Aznar, F. J., & Nadler, S. A. (2011). Erection of Ibirhynchus gen. nov.(Acanthocephala: Polymorphidae), based on molecular and morphological data. Journal of Parasitology, 97(1), 97–105.

Graham, A. (2018). The role of land bridges, ancient environments, and migrations in the assembly of the North American flora. Journal of Systematics and Evolution, 56(5), 405–429.

Grossman, J. D., & Smith, R. J. (2008). Phoretic mite discrimination among male burying beetle (Nicrophorus investigator) hosts. Annals of the Entomological Society of America, 101(1), 266–271.

Hatch, M. H. (1927). Studies on the Silphinae. Journal of the New York Entomological Society, 35, 331–370.

Hirschmann, W. (1971). Fossil mite of the genus Dendrolaelaps (Acarina, Mesostigmata, Digamasellidae) found in amber from Chiapas, Mexico. Calif Univ Publ Entomol.

Hoberg, E. P., Brooks, D. R., & Siegel-Causey, D. (1997). Host-parasite co-speciation: history, principles, and prospects. In Host–parasite evolution: General principles and avian models (pp. 212–235).

Hunter, P. E., & Rosario, R. M. T. (1988). Associations of Mesostigmata with Other Arthropods. Annual Review of Entomology, 33(1), 393–417.

Huyse, T., Poulin, R., & Théron, A. (2005). Speciation in parasites: A population genetics approach. Trends in Parasitology, 21(10), 469–475.

Hyatt, K. H. (1980). Mites of the subfamily Parasitinae (Mesostigma: Parasitidae) in the British Isles. Bulletin of the British Museum (Natural History). Zoology, 38(5), 237– 378.

Jiang, Y., Gao, M., Meng, Y., Wen, J., Ge, X.-J., & Nie, Z.-L. (2019). The importance of the North Atlantic land bridges and eastern Asia in the post-boreotropical biogeography of the Northern Hemisphere as revealed from the poison ivy genus (Toxicodendron, Anacardiaceae). Molecular Phylogenetics and Evolution, 139, 106561. https://doi.org/https://doi.org/10.1016/j.ympev.2019.106561

Kalyaanamoorthy, S., Minh, B. Q., Wong, T. K. F., Von Haeseler, A., & Jermiin, L. (2017). ModelFinder: fast model selection for accurate phylogenetic estimates. Nature Methods, 14, 587–589.

Kapli, P., Lutteropp, S., Zhang, J., Kobert, K., Pavlidis, P., Stamatakis, A., & Flouri, T. (2017). Multi-rate Poisson tree processes for single-locus species delimitation under maximum likelihood and Markov chain Monte Carlo. Bioinformatics, 33(11), 1630–1638. https://doi.org/10.1093/bioinformatics/btx025

Katoh, K., & Standley, D. M. (2013). MAFFT multiple sequence alignment software version 7: Improvements in performance and usability. Molecular Biology and Evolution, 30(4), 772–780. https://doi.org/10.1093/molbev/mst010

Keller, M. L., Howard, D. R., & Hall, C. L. (2019). Spatiotemporal niche partitioning in a specious silphid community (Coleoptera: Silphidae, Nicrophorus). Science of Nature, 106(11–12). https://doi.org/10.1007/s00114-019-1653-6

Klimov, P. B., OConnor, B. M., Chetverikov, P. E., Bolton, S. J., Pepato, A. R., Mortazavi, A. L., Tolstikov, A.V., Bauchan, G. R. and Ochoa, R. (2018). Comprehensive phylogeny of acariform mites (Acariformes) provides insights on the origin of the four-legged mites (Eriophyoidea), a long branch. Molecular Phylogenetics and Evolution, 119, 105–117.

Klompen, H., Lekveishvili, M., & Black IV, W. C. (2007). Phylogeny of parasitiform mites (Acari) based on rRNA. Molecular Phylogenetics and Evolution, 43(3), 936– 951.

Knee, W. (2017). New Macrocheles species (Acari, Mesostigmata, Macrochelidae) associated with burying beetles (Silphidae, Nicrophorus) in North America. ZooKeys, 721, 1–32. https://doi.org/10.3897/zookeys.721.21747

Knee, W., Beaulieu, F., Skevington, J. H., Kelso, S., Cognato, A. I., & Forbes, M. R. (2012). Species Boundaries and Host Range of Tortoise Mites (Uropodoidea) Phoretic on Bark Beetles (Scolytinae), Using Morphometric and Molecular Markers. PLoS ONE, 7(10), 1–15. https://doi.org/10.1371/journal.pone.0047243

Knee, W., Beaulieu, F., Skevington, J. H., Kelso, S., & Forbes, M. R. (2012). Cryptic species of mites (Uropodoidea: Uroobovella spp.) associated with burying beetles (Silphidae: Nicrophorus): The collapse of a host generalist revealed by molecular and morphological analyses. Molecular Phylogenetics and Evolution, 65(1), 276– 286.

Korallo-Vinarskaya, N. P., Krasnov, B. R., Vinarski, M. V., Shenbrot, G. I., Mouillot, D., & Poulin, R. (2009). Stability in abundance and niche breadth of gamasid mites across environmental conditions, parasite identity and host pools. Evolutionary Ecology, 23(3), 329–345.

Korn, W. (1982). Zur Fortpflanzung von Poecilochirus carabi G. und R. Canestrini 1882 (syn. P. necrophori Vitzt.) und P. austroasiaticus Vitzthum 1930 (Gamasina, Eugamasidae). SPIXIANA, 5(3), 261–288.

Kumar, S., Stecher, G., Li, M., Knyaz, C., & Tamura, K. (2018). MEGA X: molecular evolutionary genetics analysis across computing platforms. Molecular biology and evolution, 35(6), 1547–1549.

Lee, G. E., Condamine, F. L., Bechteler, J., Pérez-Escobar, O. A., Scheben, A., Schäfer-Verwimp, A., Pócs, T., & Heinrichs, J. (2020). An ancient tropical origin, dispersals via land bridges and Miocene diversification explain the subcosmopolitan disjunctions of the liverwort genus Lejeunea. Scientific Reports, 10(1), 1–13. https://doi.org/10.1038/s41598-020-71039-1

Lehmitz, R., & Decker, P. (2017). The nuclear 28S gene fragment D3 as species marker in oribatid mites (Acari, Oribatida) from German peatlands. Experimental and Applied Acarology, 71(3), 259–276. https://doi.org/10.1007/s10493-017-0126-x

Lv, J., Wu, S., Yongning, Z., Chen, Y., Feng, C., Yuan, X., Guangle, J., Junhua, D., Caixia, W., Qin, W., Lin, M., & Lin, X. (2014). Assessment of four DNA fragments (COI, 16S rDNA, ITS2, 12S rDNA) for species identification of the Ixodida (Acari: Ixodida). Parasites & Vectors, 7(1), 93.

Magalhães, S., Forbes, M. R., Skoracka, A., Osakabe, M., Chevillon, C., & McCoy, K. D. (2007). Host race formation in the Acari. Experimental and Applied Acarology, 42(4), 225–238. https://doi.org/10.1007/s10493-007-9091-0

Majka, C. G. (2011). The Silphidae (Coleoptera) of the maritime provinces of Canada. Journal of the Acadian Entomological Society, 7.

Makarova, O. L. (2013). Gamasid mites (Parasitiformes, Mesostigmata) of the European arctic and their distribution patterns. Entomological Review, 93(1), 113– 133. https://doi.org/10.1134/S0013873813010156

Mašán, P. (1999). Mites (Acarina) associated with burying and carrion beetles (Coleoptera, Silphidae) and description of Poecilochirus mrciaki sp. n. (Mesostigmata, Gamasina). Biologia, Bratislava 54, 515–524.

Matzke, N. J. (2014). Model selection in historical biogeography reveals that founder-event speciation is a crucial process in island clades. Systematic Biology, 63(6), 951– 970.

Meierhofer, I., Schwarz, H. H., & Müller, J. K. (1999). Seasonal variation in parental care, offspring development, and reproductive success in the burying beetle, Nicrophorus vespillo. Ecological Entomology, 24, 73–79.

Miao, Y., Herrmann, M., Wu, F., Yan, X., & Yang, S. (2012). What controlled Mid-Late Miocene long-term aridification in Central Asia? - Global cooling or Tibetan Plateau uplift: A review. Earth-Science Reviews, 112(3–4), 155–172.

Minh, B. Q., Nguyen, M. A. T., & Von Haeseler, A. (2013). Ultrafast approximation for phylogenetic bootstrap. Molecular Biology and Evolution, 30(5), 1188–1195.

Müller, J. K., & Eggert, A. K. (1987). Effects of carrion-independent pheromone emission by male burying beetles (Silphidae: Necrophorus). Ethology, 76(4), 297–304.

Müller, J. K., & Schwarz, H. H. (1990). Differences in carrier-preference and evidence of reproductive isolation between mites of Poecilochirus carabi (Acari, parasitidae) living phoretically on two sympatric Necrophorus species (Coleoptera, silphidae). Zoologische Jahrbücher: Abteilung Für Systematik, Geographie Und Biologie Der Tiere, 117(1), 23–30.

Navajas, M., Lagnel, J., Fauvel, G., & De Moraes, G. (1999). Sequence variation of ribosomal Internal Transcribed Spacers (ITS) in commercially important Phytoseiidae mites. Experimental and Applied Acarology, 23(11), 851–859. https://doi.org/10.1023/A:1006251220052

Navajas, M., Conte, Y. L., Solignac, M., Cros-Arteil, S., & Cornuet, J. M. (2002). The complete sequence of the mitochondrial genome of the honeybee ectoparasite mite Varroa destructor (Acari: Mesostigmata). Molecular Biology and Evolution, 19(12), 2313–2317.

Nehring, V., Müller, J. K., & Steinmetz, N. (2017). Phoretic Poecilochirus mites specialize on their burying beetle hosts. Ecology and Evolution, 7(24), 10743–10751. https://doi.org/10.1002/ece3.3591

Nehring, V., Teubner, H., & König, S. (2019). Dose-independent virulence in phoretic mites that parasitize burying beetles. International journal for parasitology, 49(10), 759–767.

Nguyen, L. T., Schmidt, H. A., Von Haeseler, A., & Minh, B. Q. (2015). IQ-TREE: A fast and effective stochastic algorithm for estimating maximum-likelihood phylogenies. Molecular Biology and Evolution, 32(1), 268–274.

Nosil, P., & Harmon, L. J. (2009). Niche dimensionality and ecological speciation. Speciation and patterns of diversity, 127–154.

Paterson, S., Vogwill, T., Buckling, A., Benmayor, R., Spiers, A. J., Thomson, N. R., Quail, M., Smith, F., Walker, D., Libberton, B., Fenton, A., Hall, N., & Brockhurst, M. (2010). Antagonistic coevolution accelerates molecular evolution. Nature, 464, 275–278.

Peck, S. B., & Anderson, R. S. (1985). Taxonomy, phylogeny and biogeography of the carrion beetles of Latin America (Coleoptera: Silphidae). Quaestiones Entomologicae, 21, 247–317.

Perotti, M. A., & Braig, H. R. (2009). Phoretic mites associated with animal and human decomposition. Experimental and Applied Acarology, 49(1–2), 85–124. https://doi.org/10.1007/s10493-009-9280-0

Ramaraju, K., & Madanlar, N. (1998). Four new Poecilochirus G. & R. Canestrini (Acarina: Parasitidae) species from Turkey. Türkiye Entomoloji Dergisi, 22(1), 3–17.

Rambaut, A., Drummond, A. J., Xie, D., Baele, G., & Suchard, M. A. (2018). Posterior summarization in Bayesian phylogenetics using Tracer 1.7. Systematic Biology, 67(5), 901–904. https://doi.org/10.1093/sysbio/syy032

Ronquist, F., Teslenko, M., Van Der Mark, P., Ayres, D. L., Darling, A., Höhna, S., Larget, B., Liu, L., Suchard, M. A., & Huelsenbeck, J. P. (2012). MrBayes 3.2: Efficient bayesian phylogenetic inference and model choice across a large model space. Systematic Biology, 61(3), 539–542.

Rueda-Ramírez, D., Santos, J. C., Sourassou, N. F., Demite, P. R., Puerta-González, A., & De Moraes, G. J. (2019). Complementary description of Africoseius lativentris and placement of Africoseius in Podocinidae (Acari, Mesostigmata) based on molecular and morphological evidences. Systematic and Applied Acarology, 24(12), 2369– 2394.

Santos, V. V. D., & Tixier, M. S. (2018). Integrative taxonomy approach for analysing evolutionary history of the tribe Euseiini Chant & McMurtry (Acari: Phytoseiidae). Systematics and Biodiversity, 16(3), 302–319.

Satou, A., Nisimura, T., & Numata, H. (2000). Reproductive competition between the burying beetle Nicrophorus quadripunctatus without phoretic mites and the blow fly Chrysomya pinguis. Entomological Science, 3(2), 265–268.

Schäffer, S., & Koblmüller, S. (2020). Unexpected diversity in the host-generalist oribatid mite Paraleius leontonychus (Oribatida, Scheloribatidae) phoretic on Palearctic bark beetles. PeerJ, 8. https://doi.org/10.7717/peerj.9710

Schedwill, P., Geiler, A. M., & Nehring, V. (2018). Rapid adaptation in phoretic mite development time. Scientific Reports, 8(1),1–10. https://doi.org/10.1038/s41598-018-34798-6

Schedwill, P., Paschkewitz, S., Teubner, H., Steinmetz, N., & Nehring, V. (2020). From the host’s point of view: Effects of variation in burying beetle brood care and brood size on the interaction with parasitic mites. PloS one, 15(1), e0228047.

Schwarz, H. H. (1996). Host range and behavioral preferences in german sibling species of the Poecilochirus carabi complex (Acari: Mesostigmata: Parasitidae). International Journal of Acarology, 22(2), 135–140.

Schwarz, H. H., & Koulianos, S. (1998). When to leave the brood chamber? Routes of dispersal in mites associated with burying beetles. Experimental and Applied Acarology, 22(11), 621–631.

Schwarz, H. H., & Müller, J. K. (1992). The dispersal behaviour of the phoretic mite Poecilochirus carabi (Mesostigmata, Parasitidae): adaptation to the breeding biology of its carrier Necrophorus vespilloides (Coleoptera, Silphidae). Oecologia, 89(4), 487–493.

Schwarz, H. H., Starrach, M., & Koulianos, S. (1998). Host specificity and permanence of associations between mesostigmatic mites (Acari: Anactinotrichida) and burying beetles (Coleoptera: Silphidae: Nicrophorus). Journal of Natural History, 32(2), 159–172.

Scott, M. P. (1998). The ecology and behavior of burying beetles. Annual Review of Entomology, 43, 595–618. https://doi.org/10.1146/annurev.ento.43.1.595

Sikes, D. S., Madge, R. B., & Newton, A. F. (2002). A catalog of Nicrophorinae (Coleoptera: Silphidae) of the world. In Zootaxa 65.

Sikes, D. S., Trumbo, S. T., & Peck, S. B. (2016). Cryptic diversity in the New World burying beetle fauna: Nicrophorus hebes Kirby; new status as a resurrected name (Coleoptera: Silphidae: Nicrophorinae). Arthropod Systematics and Phylogeny, 74(3), 299–309.

Sikes, D. S., Vamosi, S. M., Trumbo, S. T., Ricketts, M., & Venables, C. (2008). Molecular systematics and biogeography of Nicrophorus in part-The investigator species group (Coleoptera: Silphidae) using mixture model MCMC. Molecular Phylogenetics and Evolution, 48(2), 646–666.

Sikes, D. S., & Venables, C. (2013). Molecular phylogeny of the burying beetles (Coleoptera: Silphidae: Nicrophorinae). Molecular Phylogenetics and Evolution, 69(3), 552–565.

Smith, G., Trumbo, S. T., Sikes, D. S., Scott, M. P., & Smith, R. L. (2007). Host shift by the burying beetle, Nicrophorus pustulatus, a parasitoid of snake eggs. Journal of Evolutionary Biology, 20(6), 2389–2399.

Sonnenberg, R., Nolte, A., & Tautz, D. (2007). An evaluation of LSU rDNA D1-D2 sequences for their use in species identification. Frontiers in Zoology, 4(6), 1–12.

Springett, B. P. (1968). Aspects of the relationship between burying beetles, Necrophorus spp. And the mite, Poecilochirus necrophori Vitz. The Journal of Animal Ecology, 417–424.

Stadler, T. (2010). Sampling-through-time in birth-death trees. Journal of Theoretical Biology, 267(3), 396–404.

Stadler, T., Gavryushkina, A., Warnock, R. C. M., Drummond, A. J., & Heath, T. A. (2018). The fossilized birth-death model for the analysis of stratigraphic range data under different speciation modes. Journal of Theoretical Biology, 447, 41–55.

Stamatakis, A. (2014). RAxML version 8: A tool for phylogenetic analysis and post-analysis of large phylogenies. Bioinformatics, 30(9), 1312–1313.

Stefan, L. M., Gómez-Díaz, E., Mironov, S. V., González-Solís, J., & McCoy, K. D. (2018). “More than meets the eye”: Cryptic diversity and contrasting patterns of host-specificity in feather mites inhabiting seabirds. Frontiers in Ecology and Evolution, 6(JUL), 1–16.

Sun, S.-J., Horrocks, N. P. C., & Kilner, R. M. (2019). Conflict within species determines the value of a mutualism between species. Evolution Letters, 3(2), 185– 197.

Sun, S.-J., & Kilner, R. (2019). Cryptic host specialisation within Poecilochirus carabi mites explains population differences in the extent of co-adaptation with their burying beetle Nicrophorus vespilloides hosts. BioRxiv, 1223(3), 641936.

Swafford, L., & Bond, J. E. (2010). The symbiotic mites of some Appalachian Xystodesmidae (Diplopoda: Polydesmida) and the complete mitochondrial genome sequence of the mite Stylochyrus rarior (Berlese) (Acari: Mesostigmata: Ologamasidae). Invertebrate Systematics, 23(5), 445–451.

Thompson, J. N. (2009). The coevolving web of life. American Naturalist, 173(2), 125– 140. https://doi.org/10.1086/595752

Tiffney, B. H. (1985). The Eocene North Atlantic Land Bridge: Its importance in Tertiary and modern phytogeography of the Northern Hemisphere. Journal of the Arnold Arboretrum, 66, 243–274.

Wen, J., Nie, Z. L., & Ickert-Bond, S. M. (2016). Intercontinental disjunctions between eastern Asia and western North America in vascular plants highlight the biogeographic importance of the Bering land bridge from late Cretaceous to Neogene. Journal of Systematics and Evolution, 54(5), 469–490.

Wettlaufer, J. D., Burke, K. W., Schizkoske, A., Beresford, D. V., & Martin, P. R. (2018). Ecological divergence of burying beetles into the forest canopy. PeerJ, 2018(11), 1–18.

Wilson, D. S. (1982). Genetic Polymorphism for Carrier Preference in a Phoretic Mite. BioScience, 32(6), 537–537.

Wilson, D. S., & Knollenberg, W. G. (1987). Adaptive indirect effects: The fitness of burying beetles with and without their phoretic mites. Evolutionary Ecology, 1, 139– 159.

Wise, G. U., Hennessey, M. K., & Axtell, B. C. (1988). A New Species of Manure-Inhabiting Mite in the Genus Poecilochirus (Acari: Mesostigmata: Parasitidae) Predacious on House Fly Eggs and Larvae. Entomological Society of America, 81(2), 209–224.

Witaliński, W. (2000). Aclerogamasus stenocornis sp. n., a fossil mite from the Baltic amber (Acari: Gamasida: Parasitidae). Genus, 11(4), 619–626.

Xin, T., Que, S., Zou, Z., Wang, J., Li, L., & Xia, B. (2016). Complete Mitochondrial Genome of Euseius nicholsi (Ehara et Lee) (Acari: Phytoseiidae). Mitochondrial DNA Part A, 27(3), 2167–2168.

Yoder, J. B., & Nuismer, S. L. (2010). When does coevolution promote diversification? American Naturalist, 176(6), 802–817. https://doi.org/10.1086/657048

Young, M. R., Proctor, H. C., deWaard, J. R., & Hebert, P. D. N. (2019). DNA barcodes expose unexpected diversity in Canadian mites. Molecular Ecology, 28(24), 5347– 5359.

Zhang, G., Basharat, U., Matzke, N., & Franz, N. M. (2017). Model selection in statistical historical biogeography of Neotropical insects - The Exophthalmus genus complex (Curculionidae: Entiminae). Molecular Phylogenetics and Evolution, 109, 226–239.

Zhisheng, A., Kutzbatch, J. E., Prell, W. L., & Porder, S. C. (2001). Evolution of Asian monsoons and phased uplift of the Himalaya-Tibetan plate since Late Miocene times. Nature, 411, 62–66.

