## Supplementary figures and images for "Cryptic diversity within the *Poecilochirus carabi* mite species complex phoretic on *Nicrophorus burying* beetles: phylogeny, biogeography, and host specificity"

### Supplemental_Figure_1

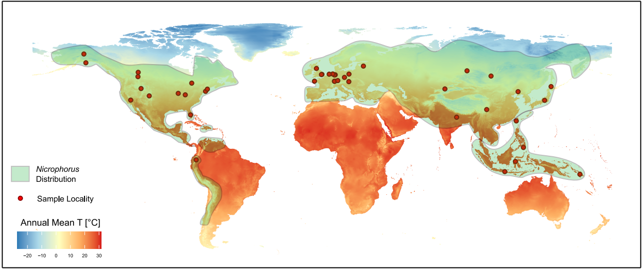

### Supplemental_Figure_2

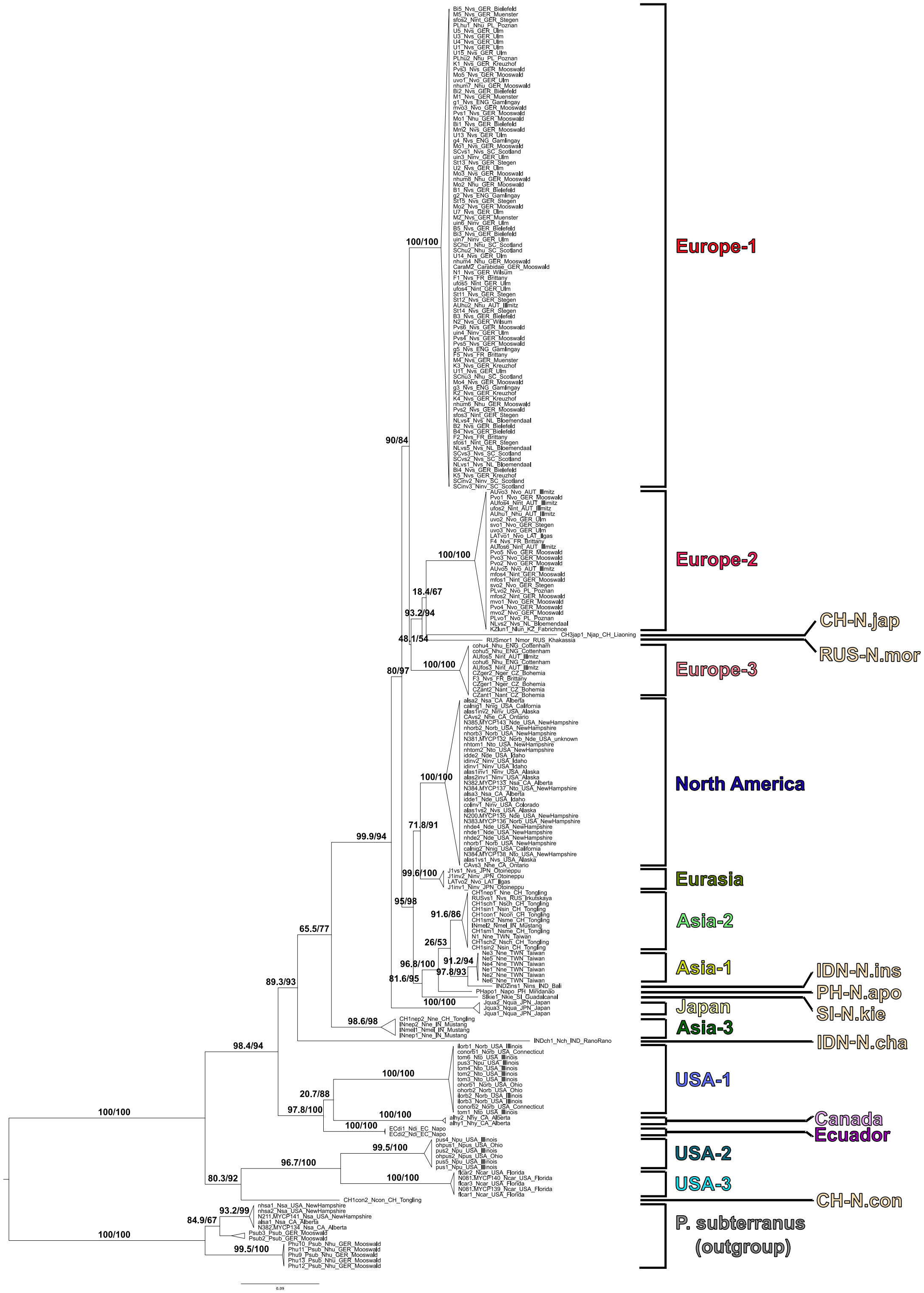

### Supplemental_Figure_3

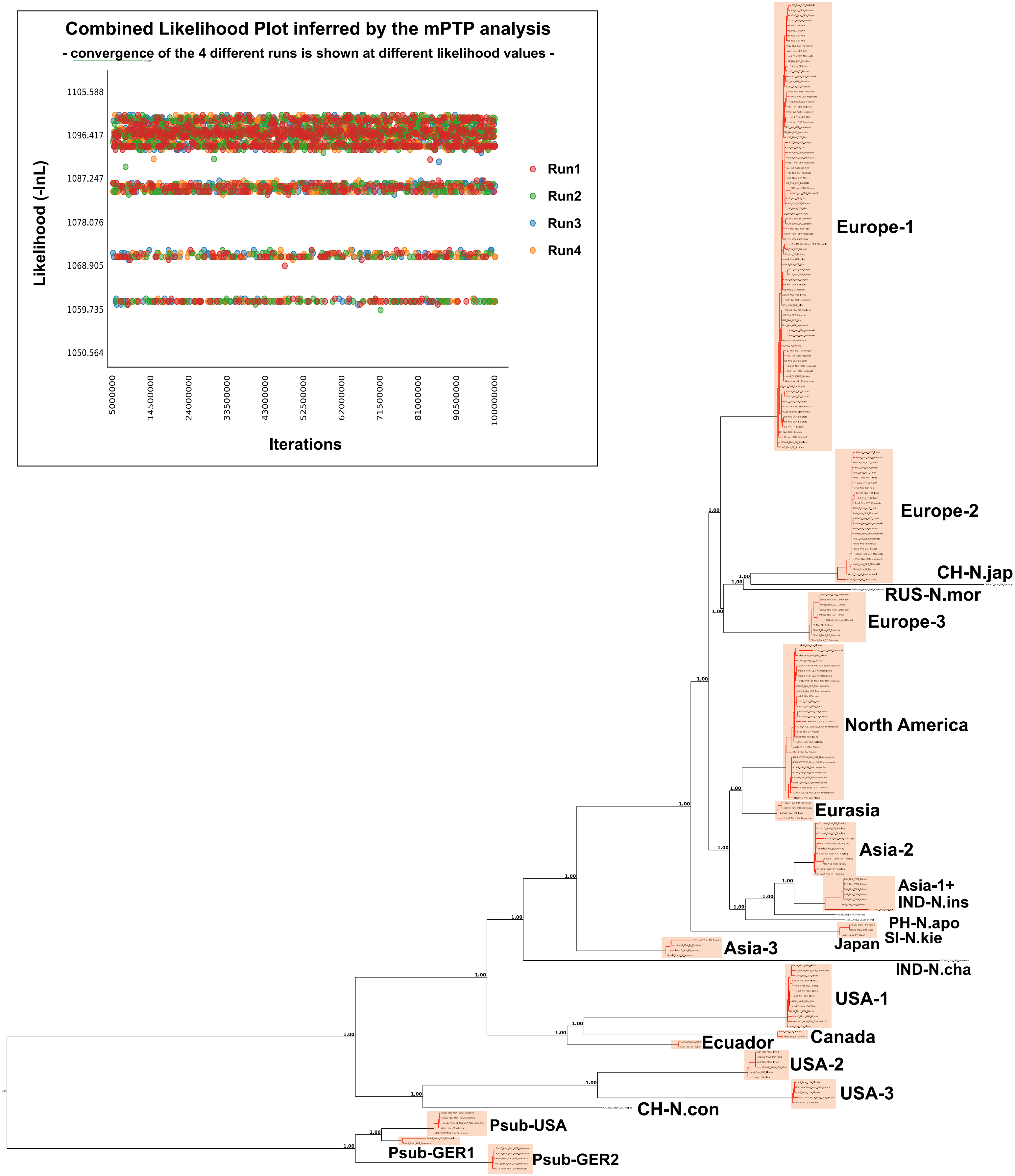

### Supplemental_Figure_4

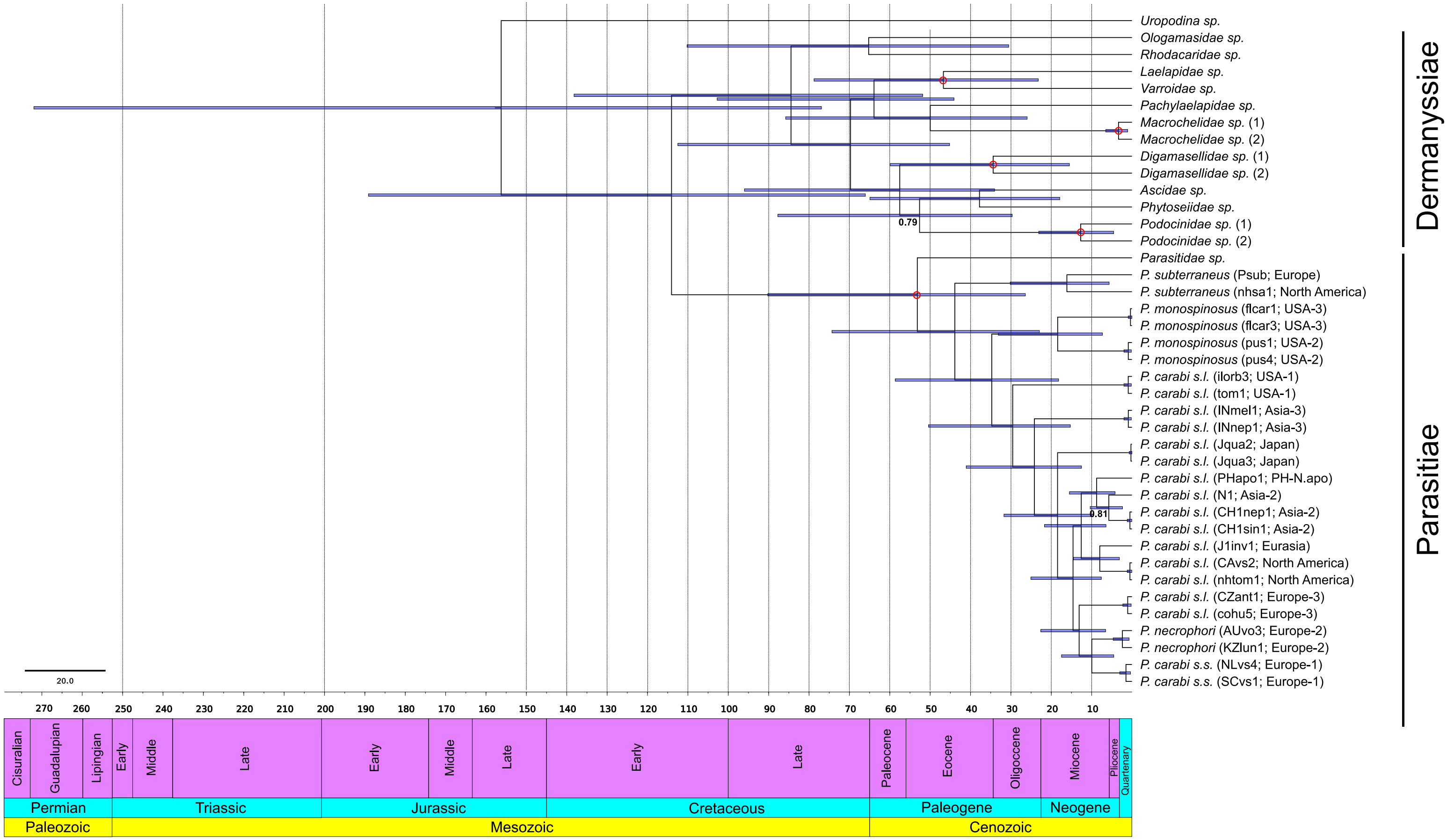

### Supplemental_Figure_5

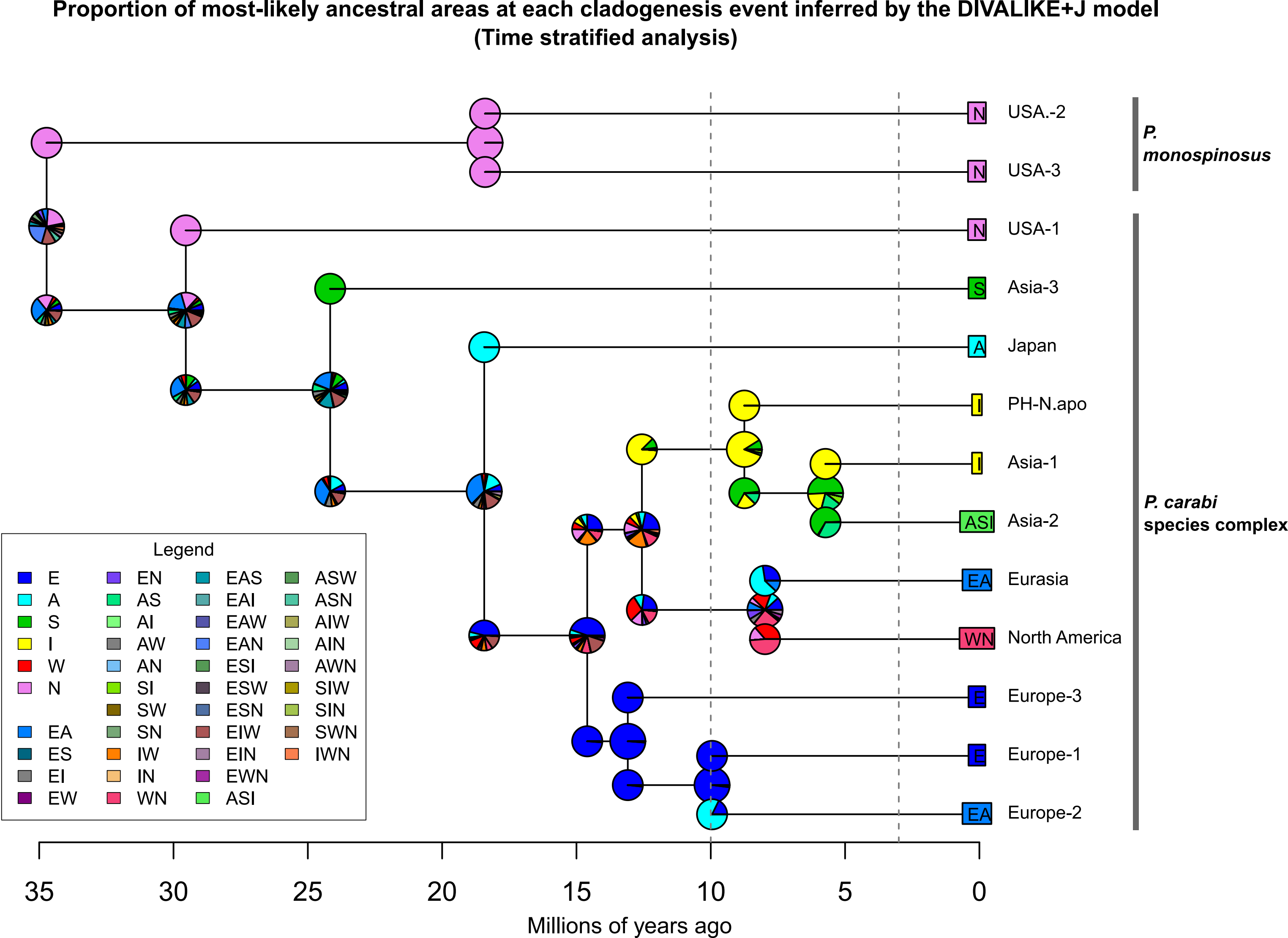
